# Distinct DNA-binding syntax and chromatin remodeling capacities of bHLH transcription factors in cell differentiation, reprogramming and cancer

**DOI:** 10.64898/2026.02.10.705010

**Authors:** Xabier de Martin, Joel Martínez-Miralles, Serafima Beletskaya, Adrià Navarro, Jan Izquierdo, Baldomero Oliva, Gabriel Santpere

## Abstract

Basic helix–loop–helix (bHLH) transcription factors orchestrate cell differentiation, reprogramming, and oncogenic transformation, yet the molecular determinants that govern their DNA-binding specificity and capacity to remodel chromatin remain incompletely understood. Here, we assemble and unify all available bHLH induction ChIP-seq datasets (74 experiments covering 17 factors) and integrate them with matched chromatin accessibility, nucleosome positioning, CpG methylation, transcriptomic profiling, methyl-HT-SELEX, and structural modeling. Using an exact hexanucleotide–based approach, we define the sequence grammar that shapes bHLH–DNA interactions and identify distinct motif architectures associated with binding to accessible versus inaccessible chromatin. CAT- and CAG-preferring bHLH factors—including proneural and myogenic regulators—display robust pioneer-like behavior, characterized by preferential engagement of closed chromatin through specific E-box variants, cooperative motif clustering, and characteristic spacing patterns. TWIST factors exhibit a unique 5-bp E-box periodicity linked to dedifferentiation programs, whereas CAC-preferring oncogenic bHLHs remain largely restricted to open chromatin. By integrating methylation and SELEX data, we reveal that CpG methylation drives a systematic shift from the canonical CAC–CAC motif toward CAT–CAC E-boxes, providing a mechanistic explanation for MYC enhancer invasion in cancer and uncovering parallel behavior in the HEY family. Nucleosome chemical mapping and structural predictions further show that nearly all bHLH factors bind preferentially at nucleosome flanks, independent of α-helix length, challenging prior models of nucleosomal engagement. Together, these results establish a unified framework linking sequence syntax, chromatin state, and transcriptional outcome across the bHLH family, providing mechanistic principles for understanding lineage specification, reprogramming, and oncogenic enhancer remodeling.

## Introduction

Basic helix-loop-helix (bHLH) transcription factors are key regulators of development and cell fate specification (*1*, *2*), controlling processes such as neurogenesis, myogenesis, and hematopoiesis. They display a broad evolutionary radiation across plants, fungi, and metazoans (*3*), and participate in a wide range of cellular and physiological functions, including responses to environmental cues, circadian rhythms, and cell cycle control (*1*, *2*).

The bHLH family of transcription factors represents a foundational genetic toolkit for developmental engineering. To date, single ectopic overexpression of bHLH transcription factors in cultured cells has proven successful in driving the differentiation of multiple cell lineages, including neurons (*4–7*), muscle cells (*8–10*), blood cells (*11*, *12*), osteoblasts (*13*, *14*), oligodendrocytes (*14–16*), melanocytes (*17*, *18*), and endothelial cells (*19*, *20*). Conversely, bHLH inductions have also proven effective in blocking cell differentiation across multiple lineages, including neurogenesis (*21–24*), myogenesis (*12*, *25*, *26*), and osteoclast (*27*), myofibroblast (*28*), and adipocyte differentiation (*29*). More strikingly, forced bHLH expression has enabled direct reprogramming between mature cell types, including conversion of astrocytes (*30–43*), fibroblasts (*10*, *44–47*), and Müller glia into neurons (*48*, *49*), as well as fibroblasts into muscle cells (*10*, *50–52*). Beyond their role in developmental reprogramming, ectopic bHLH expression has been widely used to model cell dedifferentiation (*25*, *26*) and epithelial-to-mesenchymal transition (*53–58*), processes that contribute to cancer initiation and metastasis, respectively, when dysregulated *in vivo*. Consistently, overexpression of some bHLH factors in cancer cells has been shown to enhance malignant properties (*59–63*). Crucially, beyond their practical utility in generating specific cell types, bHLH induction experiments have long served as a powerful experimental framework to dissect the molecular mechanisms underlying cell fate specification, lineage commitment, and chromatin remodeling (*5*, *9*, *10*, *50*).

While regulating such a diverse repertoire of biological processes, bHLH factors exert their action binding to a short sequence with little degeneracy, the consensus CANNTG sequence, known as the E-box, where preference for the central NN dinucleotide varies between bHLH subclasses. Each bHLH monomer binds one CAN half-site on opposite DNA strands, allowing E-boxes to be classified according to the 5′–3′ orientation of their half-sites. Based on this dimeric engagement, we employ a nomenclature previously established in our work (*64*, *65*) where motifs are defined by their constituent half-sites—labeling, for instance, the CAGATG sequences (and its reverse complement CATCTG) as CAT–CAG E-boxes.

Recent studies have demonstrated that E-box variant usage influences chromatin binding preferences and pioneer activity among bHLH transcription factors. Proneural factors such as NEUROD2 and NEUROG2 preferentially engage less accessible chromatin through the CAT-CAT and CAT-CAG E-boxes, accounting for much of the chromatin remodeling observed during neurogenesis (*64*). Earlier work by Casey et al. further showed that ASCL1, ASCL2, and MYOD1 engage closed chromatin preferentially with CAG–CAC motifs, while displaying greater association with CAG-CAG motifs in accessible regions (*9*). Despite the growing number of ChIP-seq studies following ectopic bHLH expression, systematic analyses comparing the ability of different bHLH factors to engage closed chromatin remain limited, and the molecular mechanisms underlying these differences are still poorly understood.

Here, we present the first comprehensive and integrative analysis of 74 post-induction ChIP-seq datasets covering 17 bHLH members, including representatives from all major subclasses. By combining pre- and post-induction chromatin accessibility and RNA-seq data, complemented with structural modeling, we systematically compared the capacity of bHLH factors to access closed chromatin, identified the specific E-box variants they employ, and linked these preferences to transcriptional activation outcomes. Integration of DNA methylation and methyl-SELEX data further revealed a mechanistic basis for MYC-driven enhancer invasion upon overexpression in cancer. Together, this work uncovers general principles of DNA-binding specificity within the bHLH family and defines the motif grammar underlying their competence for chromatin-remodeling and transcriptomic reprogramming.

## Results

### Collection of bHLH ChIP-seq data from induction experiments

To investigate the mechanisms of DNA binding and chromatin remodeling among bHLH transcription factors, we compiled a comprehensive catalog of 271 experiments involving single bHLH induction (**Table S1**) and systematically screened them to identify those that included ChIP-seq following TF induction. We identified 89 such studies, detailed in Table S2, and retrieved the corresponding ChIP-seq datasets from ChIP-Atlas. We used the distribution of canonical CANNTG E-box motifs around ChIP-seq peak summits as a proxy for ChIP-seq data quality (**Fig. S1A**). Based on this analysis, we retained 74 datasets corresponding to 17 bHLH TFs that showed significant E-box centrality compared to a uniform distribution. These factors span all major subgroups of the bHLH family, including: proneural TFs (NEUROG2, NEUROD1, NEUROD2, ASCL1, OLIG2); neural crest cells, mesodermal and mesenchymal regulators (MITF, TWIST1, TWIST2, MESP1, MSGN1); myogenic factors (MYOD1, MYF5); proto-oncogenes (MYC, MYCN); transcriptional repressors (HEY1, HEY2, BHLHE40); and the broadly expressed E-protein TCF4.

### Global classification of bHLH binding preferences

The 74 induction experiments spanned diverse cell types, enabling the identification of general principles of bHLH DNA binding. According to their intrinsic *in vitro* binding preferences, bHLH proteins cluster into three classes defined by their half-site specificity: CAT (e.g., NEUROG2, NEUROD2, TWIST1), CAG (e.g., MYOD1, ASCL1, TCF4), or CAC (e.g., MYC, HEY1, MITF) (*65*, *66*). At endogenous expression levels, intrinsic DNA-binding preferences may be obscured by interactions with cellular co-factors and co-expressed partners. Although overexpression-based experiments may generate non-physiological binding profiles, they substantially increase the power to detect subclass-specific binding preferences. Consistent with this, we observed that the binding landscape of bHLH factors is markedly expanded after their forced overexpression (**Fig. S1B**), and that this expansion is mediated by binding to their cognate E-box motifs (**Fig. S1C**). This expansion was most pronounced for factors that are normally absent or expressed at very low levels in the induced cell types, whereas for CAC-preferring factors it more closely reflects a classical overexpression scenario. Notably, this expansion differentially impacts genomic targeting, particularly for CAC-preferring factors, with MYC consistently increasing the proportion of enhancer vs promoter binding (**Fig. S1D**).

### E-box usage and motif centrality across bHLH subfamilies

We quantified and analyzed the distribution of each E-box variant around the summits of ChIP-seq peaks across all induction experiments (**Fig. 1A, Fig. S2**). Among factors in the CAT-preferring cluster, CAT-CAG emerged as the most abundant and centrally enriched motif (*65*), followed by CAG-CAG. Although the palindromic CAT-CAT motif is less frequent overall, it shows the third highest degree of central enrichment, consistent with observed *in vivo* binding patterns (*64*). This strong central enrichment is specifically prominent in the CAT cluster, compared with any other bHLH factors analyzed (**Fig. 1A, Fig. S2**). CAG-preferring factors showed strong enrichment for palindromic CAG–CAG E-boxes, followed by CAG–CAC variants. An exception was the E-protein TCF4, which displayed preferential enrichment for CAG–CAC rather than CAG–CAG E-boxes (**Fig. 1A, Fig. S2**).

**Fig. 1.**
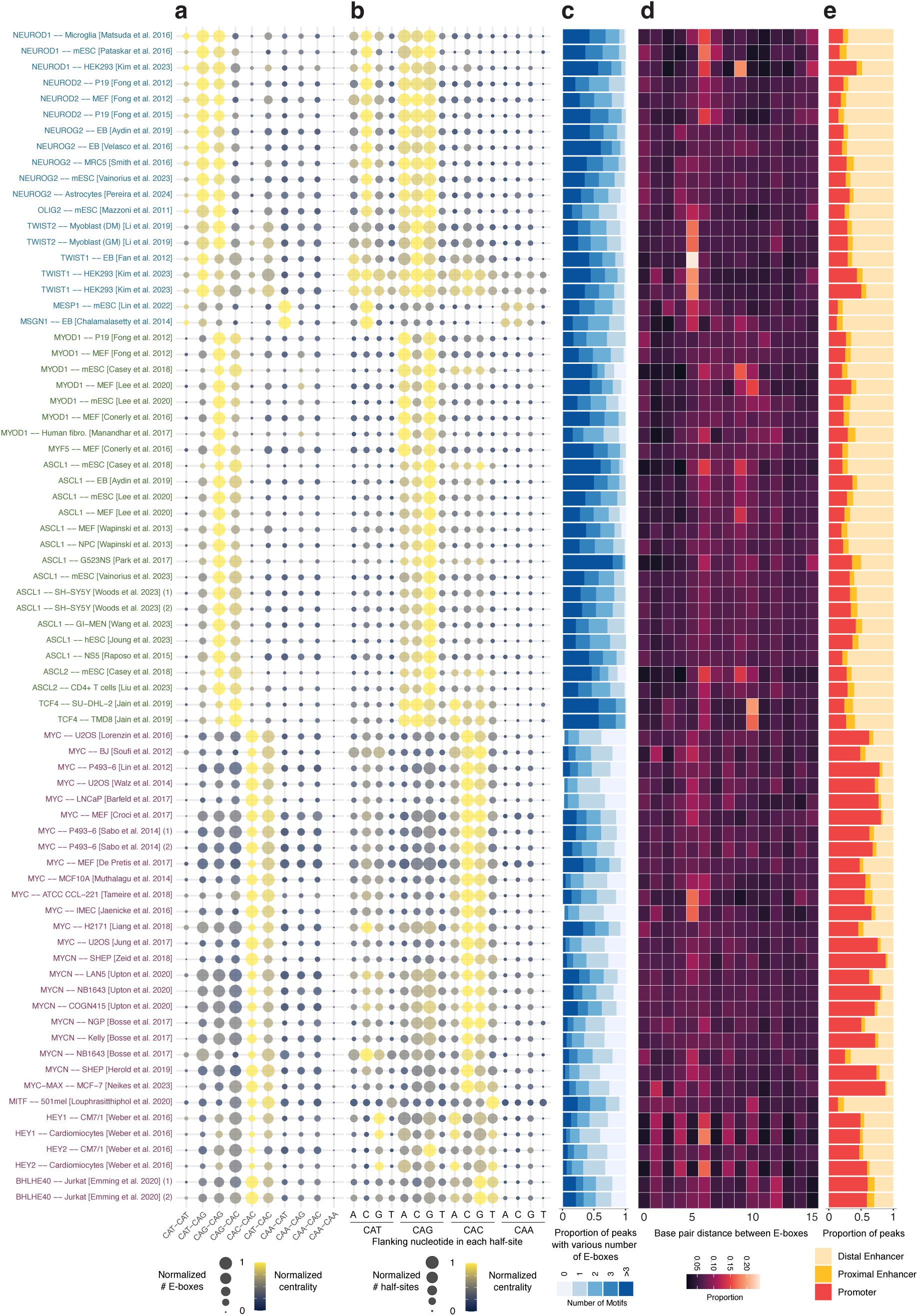
Regulatory syntax of E-box motifs across bHLH induction ChIP-seq datasets. **A**) Balloon plot showing the enrichment of E-box variants in each ChIP-seq experiment. The number of motifs per peak is depicted with the size of the balloon, and the centrality, with the color scale. Both metrics are min-max normalized by experiment. B) Balloon plot depicting the enrichment of individual E-box half-sites extended by one flanking nucleotide (NCAN). Half-sites from each E-box were analyzed independently and treated as distinct motifs; balloon size and color encode motif abundance per peak and positional centrality, respectively. **C**) Stacked barplot displaying the proportions of peaks encompassing different numbers of E-box motifs per peak. **D**) Heatmap showing the proportions of base-pair distances between adjacent canonical E-boxes located within peaks. **E)** Genomic distribution of peaks relative to the Transcription Start Site (TSS). Regions are classified as proximal promoters (< ±2 kb), proximal enhancers (±2–4 kb), and distal enhancers (> ±4 kb).

The CAC-CAC palindrome is the preferred motif for the CAC-preferring bHLH subclass, including MYC, MYCN, and HEY factors, followed by CAT-CAC (**Fig. 1A**). Interestingly, CAT-CAC motifs are also frequently found in the binding sites of TWIST factors. While the CAA-CAA palindrome remains the only E-box type not bound by any bHLH dimer—consistent with previous reports (*67*) —the CAA half-site can nonetheless be recognized when paired with a preferred partner site. For example, we observed a slight central enrichment of the heterotypic CAA–CAG motif in MyoD1 datasets (Fig. 1A). This is consistent with observations of the *C. elegans* MRF ortholog, HLH-1, which also targets the CAA-CAG E-box (*68*). Another compelling example is provided by Mesp1 and Msgn1, which display a clear preference for the CAA–CAT motif, a binding profile that diverges markedly from that of their closest phylogenetic relatives within the CAT cluster (*65*). Notably, these are the only factors within this cluster known to be involved in early mesoderm development, suggesting a potential E-box specialization linked to this early developmental process.

### Flanking nucleotide preferences around E-box half-sites

Previous structural studies revealed contacts between bHLH factors and nucleotides flanking the E-box (*67*, *69*). Such contacts may impose preferences for bases outside the core motif, as reported for specific family members, and confer binding specificity (*70–75*). For example, NEUROD2 and NEUROG2 show a preference for a C preceding CAT half-sites and a G preceding CAG half-sites (*64*). Whether these biases are intrinsic to particular half-sites across the broader bHLH family has remained unclear. To address this, we quantified the enrichment of flanking nucleotides associated with each half-site for every bHLH factor. We found that each half-site was associated with characteristic nucleotide preferences, a general feature shared by most—but not all—members of each bHLH subfamily. Specifically, we confirmed the enrichment of a C preceding CAT half-sites in NEUROD2 and NEUROG2 across multiple datasets and extended this observation to other CAT-preferring factors, including NEUROD1 and OLIG2, with the exception of TWIST proteins. CAG-preferring factors also displayed systematic variation: MYOD1 and MYF5 favored ACAG and GCAG half-sites, whereas ASCL1 was more selectively enriched for GCAG-containing E-boxes, and TCF4 preferred ACAG and CCAG. These findings suggest that flanking dinucleotides contribute to binding specificity even among bHLH proteins recognizing the same E-box variant. Finally, MYC and MYCN consistently showed enrichment for a C or G preceding their preferred CAC half-site. Strikingly, nearly all factors disfavored a T in this position, consistent with in vitro observations for TAL1/TCF3 heterodimers (*72*), TCF3 (*76*) or MYC (*77*, *78*), with the notable exception of MITF, which preferentially bound “TCAC” half-sites, as has been observed previously in melanocytes and described as a discriminative feature between MITF and MYC-MAX binding (*79*). In summary, the half-site preferences of individual bHLH monomers give rise to a distinct, half-site–specific flanking pattern through their dimeric engagement of E-box motifs.

### Binding to degenerate E-box variants

We investigated whether bHLH factors can bind degenerate E-box variants in a manner dependent on their preferred motif type. To do so, we analyzed the distribution around the ChIP-seq summit of E-boxes containing a single nucleotide mutation in the core ‘CA’ dinucleotide in one of the half-sites. We found that tolerance to these mutations varied by E-box (**Fig. S2**). For instance, both CAT- and CAG-preferring factors showed modest central enrichment for degenerate CAG-CAG motifs, though significantly lower than for their canonical counterparts. MYC and MYCN also displayed some tolerance, particularly toward mutated CAC-CAC motifs. Interestingly, the repressor factors HEY1 and HEY2 exhibited even higher centrality for degenerate motifs such as CAC-CAC and CAT-CAC than for their canonical versions (**Fig. S2**). This observation is consistent with the ability of HEY factors to heterodimerize with HES factors where one subunit binds a CAC half-site and the other targets a non-canonical sequence such as CTN or CGC (*74*).

### Motif clustering, spacing and local cooperativity

The presence of multiple E-box motifs in proximity within regulatory regions has been implicated in strengthening both the binding efficiency of bHLH transcription factors and the activation of downstream genes (*50*, *80–82*). To explore this, we analyzed the clustering of E-box motifs in ChIP-seq peaks among induction experiments. Compared to E-box occurrences in size-matched peak flanking regions (**Fig. S3A)**, genomic regions bound by bHLH transcription factors show a higher frequency of clustered E-box motifs, particularly for factors that prefer CAG and CAT half-sites (**Fig. 1C**). In contrast, CAC-preferring factors tend to exhibit fewer E-box motifs per peak. This is consistent with previous findings showing that MYC binds to approximately half of euchromatic regions in the absence of E-box motifs (*83*), with its binding pattern correlating more strongly with RNA polymerase II occupancy than with the presence of E-boxes (*71*).

We next investigated whether clustered E-boxes exhibit a specific spacing pattern (**Fig. 1C**). TWIST1 and TWIST2 showed the strongest enrichment for tandem E-boxes separated by 5bp. This spacing is consistent with previous reports indicating that TWIST1 preferentially binds DNA as a tetramer, formed by two TWIST1/E-protein heterodimers, which is required for TWIST1-mediated induction of the epithelial-mesenchymal transition (*84*). We did not observe this specific spacing pattern in any other bHLH member. Instead, other subfamilies exhibit distinct preferences; for instance, NEUROD family members appear to preferentially bind E-box pairs separated by 6bp, consistent with *in vivo* observations for NEUROD2 (*64*). ASCL1 and MYOD1 exhibit a preference for E-box pairs spaced 6 and 9bp apart in datasets generated by Casey et al. (2018) in embryonic stem cells (ESCs) (*9*) and a 10bp spacing pattern in datasets from Lee et al. (2023) using mouse embryonic fibroblasts (MEFs) (*10*). We observed a similar 10bp-spacing enrichment in two independent TCF4 ChIP-seq datasets derived from distinct cell lines (**Fig. 1C**). No spacing pattern was detected in random sets of sequences with matched dinucleotide composition (**Fig. S2B**).

To assess whether E-boxes were the sole class of motifs enriched at peak summits, we calculated the central enrichment of 286 archetypal transcription factor motifs from Vierstra et al. (2020) (*85*). After excluding motifs that contain E-box–like sequences, we found no evidence of secondary enrichment for any other archetypal motif (**Fig. S1C**). These results suggest that bHLH factors bind DNA independently, without a consistent requirement for co-binding with non-bHLH transcription factors.

Finally, we examined the genomic context of these binding events and found that CAT- and CAG-preferring bHLH factors predominantly bind to distal enhancers, whereas CAC-preferring bHLH factors are more frequently associated with promoter regions, an exception being MITF, which displays a more enhancer-like binding profile (**Fig. 1D**).

Taken together, these analyses define general principles of bHLH binding across different subfamilies, confirming previous observations for individual factors and extending them to additional members within each class. Differences in E-box usage, motif density, spacing and genomic distribution emerge as intrinsic properties of each transcription factor, largely independent of the cellular context.

### DNA binding features vary with chromatin accessibility

A growing body of evidence indicates that members of the bHLH family differ in their ability to bind closed chromatin and initiate regulatory programs, both in their native cellular context and when ectopically expressed. Having established the general binding features of each bHLH factor in the previous analyses, we next investigated how these properties correlate with the chromatin accessibility landscape of the cellular populations prior to factor induction.

For each cell type in each dataset, we obtained ATAC-seq or H3K27ac ChIP-seq data (**Table S4**) and quantified the maximum signal overlapping a window around the summit of each bHLH ChIP-seq peak. Peaks were then ranked and partitioned into quartiles based on their pre-induction accessibility levels. This analysis revealed that CAT- and CAG-preferring bHLH factors were capable of binding regions across a broad range of chromatin accessibility, including low-accessibility sites. In contrast, except for MITF, all CAC-preferring bHLH factors, such as MYC, MYCN, and the HEY1/2, were predominantly restricted to highly accessible chromatin regions across all evaluated cell types (**Fig. 2A**).

**Fig. 2.**
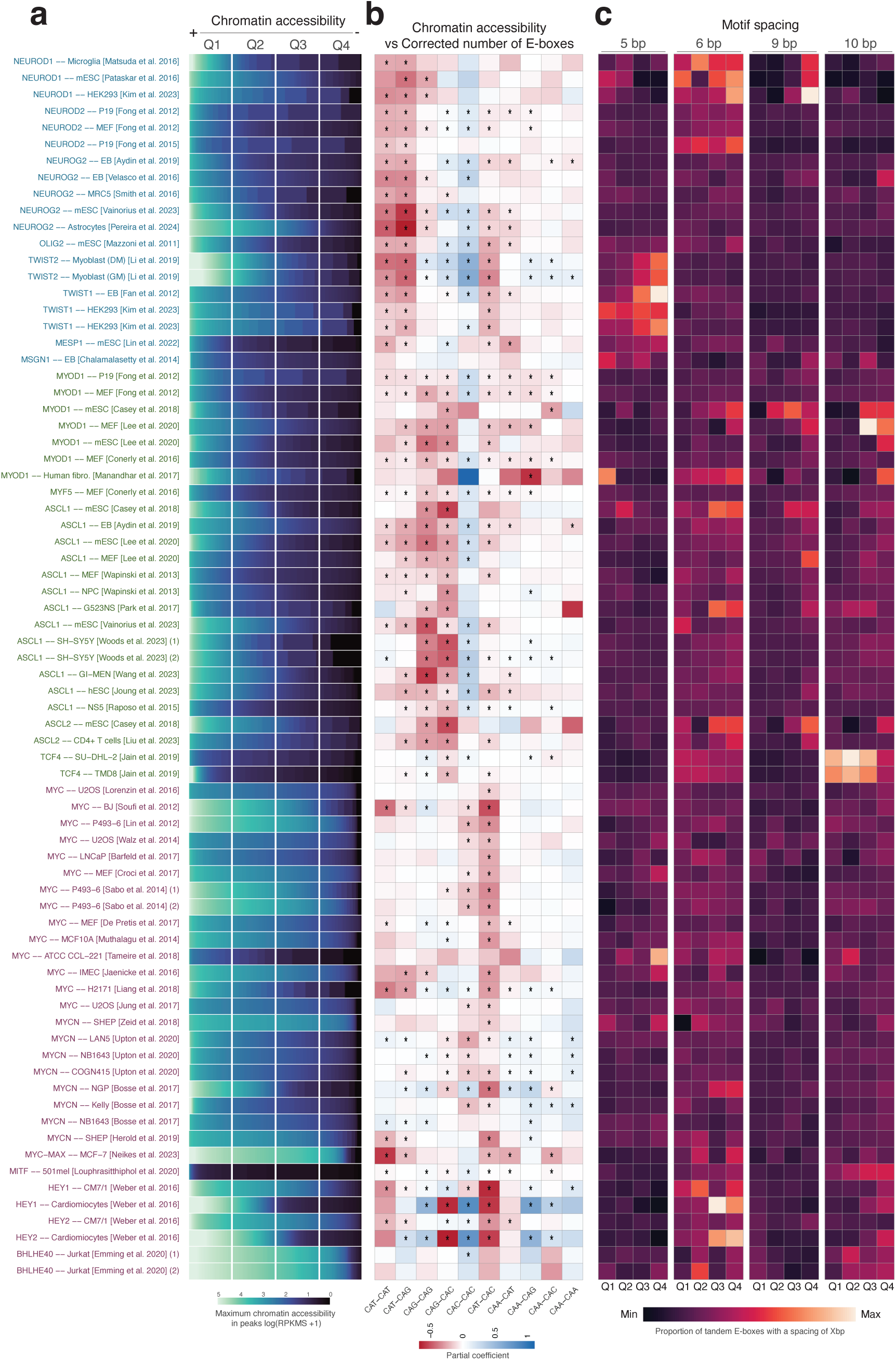
Characterization of DNA-binding grammar of bHLH factors in the pre-induction chromatin landscape. **A**) Heatmap displaying pre-induction accessibility across ChIP-seq peaks for each study. Peaks were sorted by maximum accessibility within a 100bp window around their summit(log-transformed RPKM + 1) and partitioned into 271 equal-sized bins (the minimum number of peaks found across all datasets) to allow cross-study comparison. Each cell represents the mean accessibility of the peaks within that bin. **B**) Heatmap displaying the partial coefficients derived from a multivariate linear model predicting pre-induction chromatin accessibility of the bound region (log-transformed RPKM + 1) based on the enrichment of all E-box variants within ChIP-seq peaks. Negative coefficients (red) indicate that the motif is more enriched the less accessible was the region, whereas positive coefficients (blue) indicate enrichment in regions with higher basal accessibility. Asterisks denote statistical significance after Benjamini-Hochberg correction (q < 0.01). **C)** Heatmaps showing pacing syntax across chromatin states. The value in each cell represents the proportion of tandem E-boxes separated by a specific distance (5,6,9 and 10bp for each heatmap) as compared to the other tandem E-boxes. Each heatmap is normalized relative to its maximum..

Previous observations suggest that both the number of clustered E-box motifs and their specific type influence the ability of bHLH factors to bind low-accessibility chromatin. However, this association has rarely been investigated in induction experiments, i.e., only in three out of the 74 experiments (**Table S5**). To quantify this relationship, we examined motif-specific enrichment in ChIP-seq peaks corrected by E-box density in size-matched flanking regions and related it to pre-induction chromatin accessibility of the peak (**Fig. 2B, Fig. S4A**). Notably, motif effects were estimated using a multivariate regression framework where pre-induction chromatin accessibility was modeled as a function of all E-box subtypes simultaneously, thereby isolating the contribution of each specific motif while controlling for the presence of other E-box types at the bound site. This analysis showed that CAT-preferring factors mainly use CAT-CAT and CAT-CAG motifs when binding to closed chromatin. TWIST1 and TWIST2 additionally exhibited significant enrichment of the CAT–CAC motif in closed chromatin, whereas for the mesodermal factor MESP1 the CAT–CAA motif was the most strongly associated with less accessible regions (**Fig. 2B, Fig. S4A**). By contrast, the CAG–CAG motif, despite its high abundance and strong centrality in this bHLH subclass, displayed a markedly weaker association with chromatin accessibility. For factors in the CAG subclass, both preferred motifs, CAG–CAG and CAG–CAC, showed consistent associations with lower-accessibility chromatin. Finally, among CAC-preferring factors, MYC displayed a striking deviation from the canonical pattern, with CAT–CAC emerging as the motif most strongly associated with less accessible chromatin. By contrast, MYCN and HEY factors showed additional associations with CAC–CAC and CAG–CAG motifs, respectively, alongside CAT–CAG.

We found that E-box spacing patterns were strongly influenced by chromatin accessibility among the bHLH factors that displayed such features. For example, the TWIST-specific 5 bp spacing was more prominent in peaks overlapping inaccessible chromatin (**Fig. 2C, Fig S4C**). Likewise, the 6 bp and 9–10 bp spacings observed in subsets of ASCL1 and MYOD1 datasets were also more pronounced in less accessible regions. Interestingly, although NEUROD1 globally favored 6 bp spacing, it showed a relative enrichment for 9 bp spacing specifically in chromatin that was initially closed. These results indicate that recurrent spacing preferences among CAT- and CAG-preferring bHLH factors are preferentially manifested in inaccessible chromatin, suggesting that higher-order E-box architecture may facilitate binding in compacted regulatory regions.

Collectively, these results show that bHLH factors vary widely in their ability to engage closed chromatin, and that this capacity is determined by specific sequence features—namely motif density, E-box subtype, and spacing. CAT- and CAG-preferring factors show a consistent ability to bind to closed chromatin, mediated by subclass-specific binding patterns. Paradoxically, while the binding of CAC-preferring factors is mainly restricted to accessible chromatin, these factors show a strong E-box signature associated with variation in pre-induction chromatin accessibility. And even more intriguingly, this signature does not point to the preferred and widely studied CAC-CAC, but for the less-optimal CAT-CAC variant.

### bHLH interaction with methylated DNA

Chromatin regions with low accessibility often coincide with high levels of CpG methylation, which can directly affect transcription factor binding. Because certain E-box variants contain CpG sites—specifically the canonical CACGTG (CAC–CAC) motif—we examined how CpG methylation influences bHLH factor binding across different chromatin contexts. MYC is known to invade enhancers upon overexpression in cancer, and enhancers show a generally higher and more dynamic methylation status. We hypothesize that MYC and other CAC-preferring factors could be redirected to other E-boxes in highly methylated regions, since CpG methylation could impair the recognition of their canonical CACGTG motif.

To test this hypothesis, we first reanalyzed HT-SELEX and methyl-HT-SELEX datasets for the CAC–CAC–preferring bHLH members included in our study: MAX (the obligate *in vivo* MYC partner), HEY1, and HEY2. In unmethylated conditions, the CpG-containing CAC–CAC motif was the most strongly enriched. In contrast, under DNA methylation, CAT–CAC motifs became predominant, indicating a clear methylation-dependent shift in E-box usage (**Fig. 3A**).

**Fig. 3.**
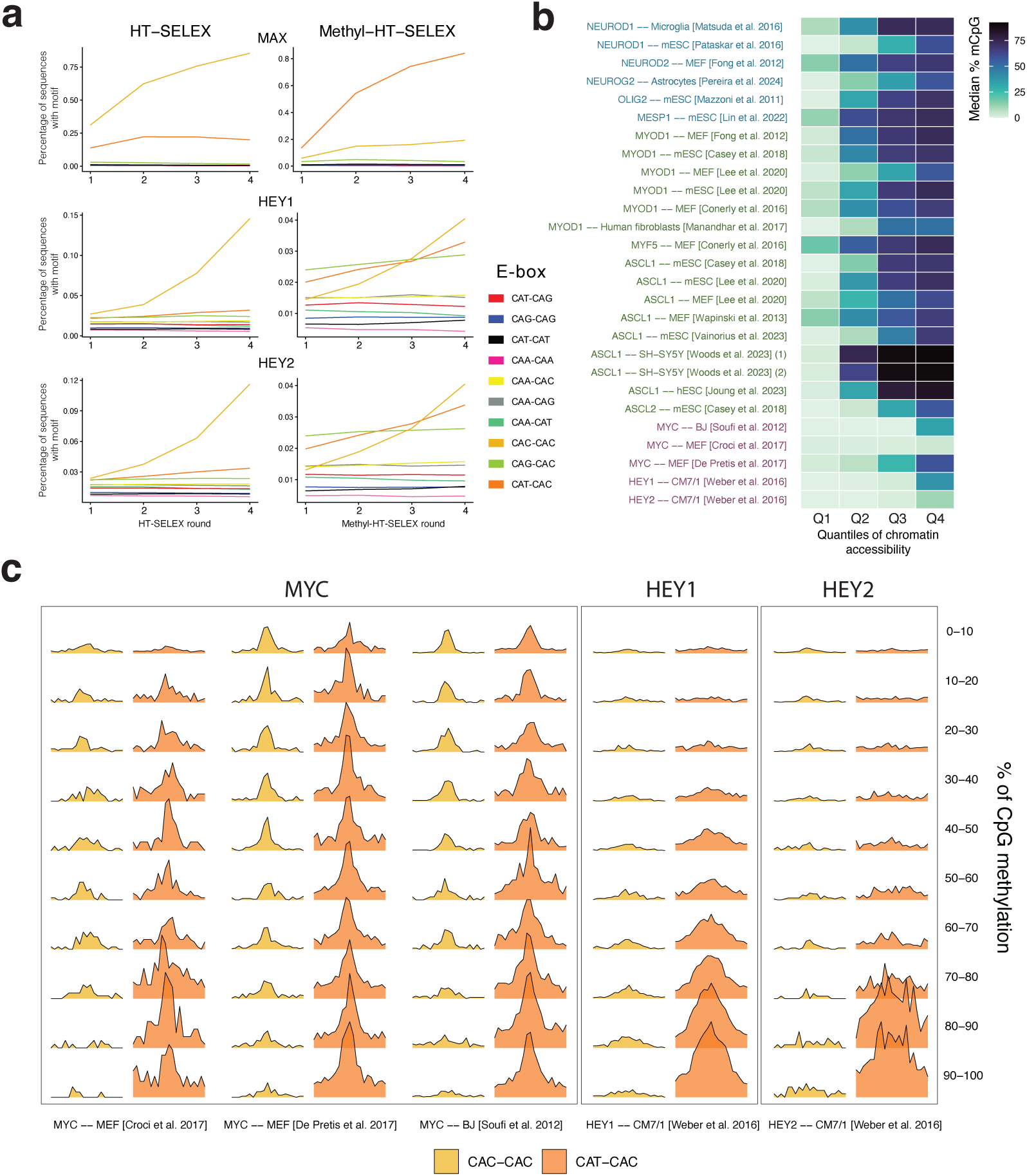
Variation in DNA-binding grammar as a function of pre-induction CpG methylation. **A**) Lineplots where the y-axis represents the proportion of sequences containing each type of E-boxes in the consecutive rounds of HT-SELEX and Methyl-HT-SELEX (X axis). **B**) Heatmap displaying the variation of CpG methylation levels in pre-induction chromatin accessibility quartiles (Q1: highest; Q4: lowest). Methylation was calculated as the mean percentage of methylated CpGs per individual peak and then averaged across the peaks of each accessibility quartile. **C**) Frequencies of motifs per peak of CAC-CAC and CAT-CAC variants in a window of 400bp around the summits of peaks in the 4th quartile of chromatin accessibility (closed chromatin) and further stratified in bins of %CpG methylation, for MYC and HEY1/HEY2 studies. Motif frequencies per peak are normalized relative to the maximum global value in each study, to highlight the relative change of a motif’s enrichment across methylation bins, and to be able to compare the net effect between the two variants in each bin.

We next set out to determine whether this methylation-dependent shift was also evident following ectopic TF overexpression in cell culture. We leveraged cell-type-matched DNA methylation data from 27 of the 76 induction experiments, including five involving MYC, HEY1, and HEY2. For each transcription factor and pre-induction chromatin accessibility quartile, we averaged the %methyl-CpG values of the CpGs that fell within the ChIP-seq peaks (**Table S6**). As expected, there was an overall increase in CpG methylation levels in less accessible chromatin. CAT- and CAG-preferring factors frequently bound these highly methylated regions. In contrast, and consistent with their preference for the CpG-containing E-box motif, MYC and HEY factors were predominantly associated with regions of lower CpG methylation compared to CAT- and CAG-preferring bHLHs (**Fig. 3B**), with a minority of their peaks overlapping highly methylated CpGs in the 4th quartile (least accessible chromatin) (**Fig. S5A**). To understand how MYC and HEY interact with these highly methylated regions, we binned the 4th quartile by methylation levels and examined the E-box usage in these bins. Interestingly, CAT-CAC E-boxes showed a clear enrichment commensurate with increasing methylation, at the expense of the canonical CAC–CAC motif. No other methylation-associated shifts in E-box usage were detected among the remaining bHLH factors (**Fig. S5B**). These results indicate that DNA methylation redirects MYC and HEY from canonical CAC–CAC sites toward atypical CAT–CAC variants. This flexibility in motif selection enables these factors to invade hypermethylated, restrictive enhancers when overexpressed, effectively expanding their regulatory reach.

### bHLH interaction with nucleosomes

Although many bHLH factors can bind to closed chromatin to some extent, the mechanisms underlying this ability remain poorly understood, particularly whether such associations involve direct interactions with nucleosomes. To address this, we used high-resolution nucleosome positioning maps from mouse ESC (mESC) generated by chemical mapping (*86*), which provide base-pair–level localization of nucleosome dyads and are minimally affected by AT-content bias (*86*, *87*). For all 17 bHLH induction studies performed in mESCs, we computed the distance between each ChIP-seq summit and the nearest nucleosome dyad and quantified the distribution of summits across defined distance intervals. Most bHLH factors displayed a depletion of summits near the dyad center, indicating a preference for binding at linker regions (**Fig. 4A**). In contrast, when we performed the same analysis for two well-characterized pioneer factors, OCT4 and SOX2, we observed a relative enrichment of summits around nucleosome dyads. These results suggest that, despite the reported pioneer activity of several bHLH members such as NEUROD, NEUROG, and ASCL1, they primarily engage nucleosome-bound DNA by binding within linker regions.

**Fig. 4.**
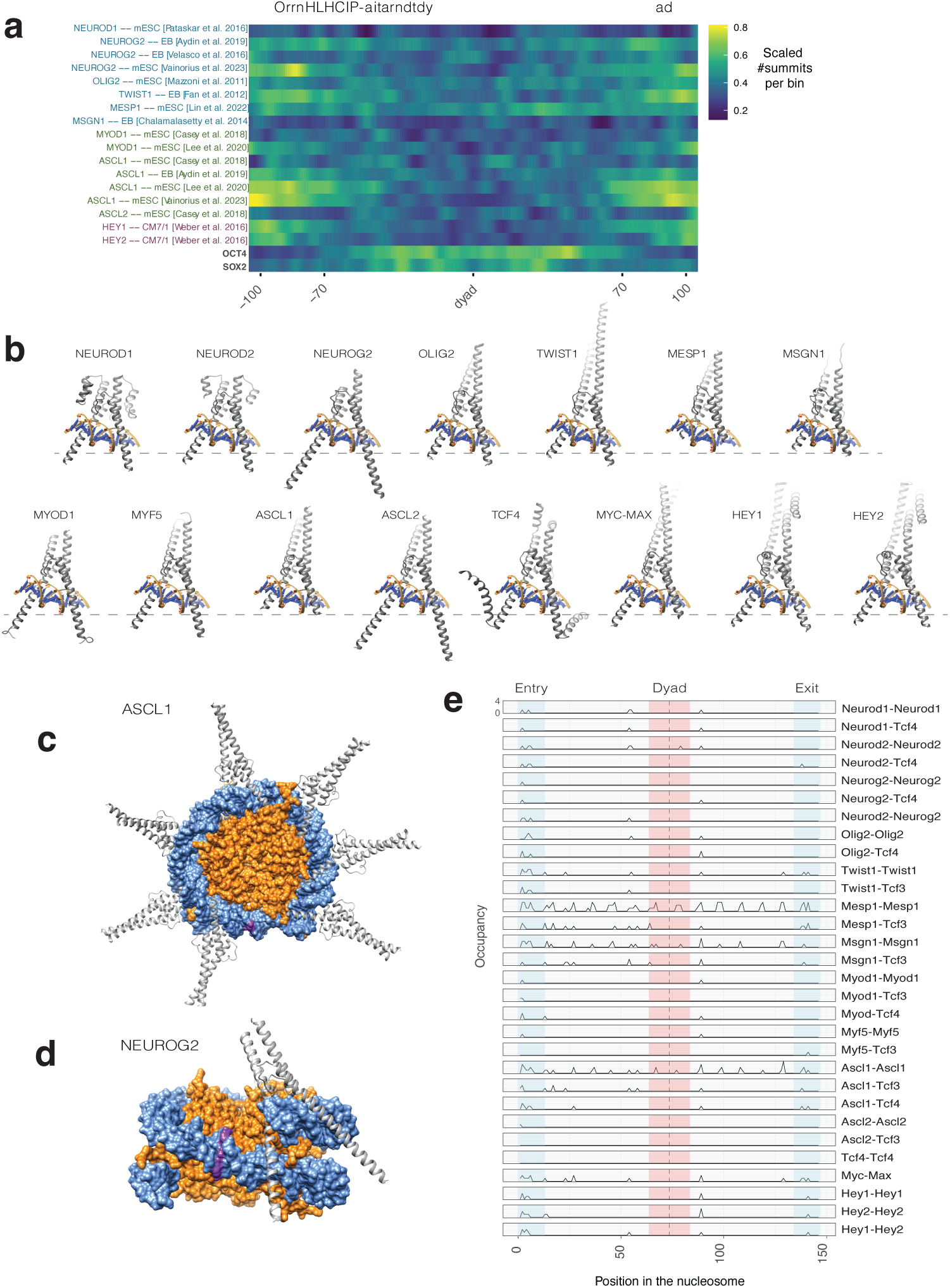
Positional mapping and steric feasibility of TF-nucleosome co-occupancy. **A**) Heatmap displaying the density of ChIP-seq summits relative to the center of the nearest nucleosome dyad (+-100bp). Summit frequency values are computed by 1bp bins around the dyad, scaled by study and then Gaussian-smoothed, illustrating the varying degrees of nucleosome-TF co-occupancy or displacement across different TFs. **B**) Structural models of bHLH dimers were generated using AlphaFold3 with DNA sequences containing an E-box motif. Models were visualized in Chimera and trimmed to remove unstructured terminal regions, retaining only well-defined and contiguous secondary-structure elements corresponding to the DNA-binding domains of the transcription factors. **C-D**) Structural models illustrating contacts of ASCL1 and NEUROG2 with nucleosomal DNA. **E**) Line plot displaying the number of transcription-factor occupancies along the nucleosomal DNA quantified by sliding the E-box window from positions 1 to 146. A maximum of two steric clashes per model was permitted.

A previous study comparing the basic helix length of six bHLH factors suggested that a shorter helix may promote more effective interactions with nucleosomal DNA. To investigate this in greater detail, we used AlphaFold3 to predict the structures of a selection of 15 bHLH factors (**Fig. 4B**). Consistent with earlier reports, ASCL1 exhibited a short basic α-helix; this feature was also observed in MSGN1, with MESP1 displaying an even shorter helix. In contrast, most CAG-preferring factors, HEY proteins and CAT-preferring bHLH factors generally harbored long helices—most notably NEUROG2 and ASCL2. Interestingly, TCF4 presented a unique additional α-helix emerging from a short loop at the end of the basic domain.

We next applied a new structural modeling approach, ModCRE, to estimate the potential nucleosome occupancy of each of the 15 bHLH AlphaFold3 structures, using a DNA template in which an E-box was positioned at every possible location, without steric clashes at 4 Å (**Fig. 4C**). Factors with short helices, such as ASCL1, established multiple contacts with the nucleosome, showing a periodic pattern in which the E-box was accessible on the outer surface of the DNA. In contrast, NEUROG2—with its long helix—could bind only at DNA ends adjacent to the linker. Systematic analysis of contact frequency at each nucleosomal position revealed a strong association between α-helix length and nucleosome interaction mode. Only ASCL1, MESP1, and MSGN1 could engage in this periodic binding, whereas factors with super-long helices bound exclusively at the less tightly wrapped DNA ends, consistent with the previously described “breathing mode” (*88*). Most bHLH factors with intermediate helix lengths could bind both through the breathing mode and at specific positions flanking the dyad. Notably, MYC–MAX has been experimentally shown to prefer end binding (*89*). TCF4 homodimers were unable to bind the nucleosome at any position without steric clashes, consistent with their inability to engage closed chromatin (**Fig. 4C**).

In summary, nucleosome mapping analyses show that bHLH factors predominantly bind at nucleosome flanks rather than the dyad. Remarkably, this bias persists even in short-helix bHLHs, for which our structural analysis indicates that periodic nucleosomal binding would be sterically possible, suggesting that additional constraints limit their engagement with nucleosomal DNA.

### Global chromatin-remodeling capacity across bHLH subclasses

Having characterized how bHLH factors engage DNA across varying chromatin accessibility and CpG methylation levels, we next examined their chromatin-remodeling capacities by comparing epigenomic data before and after factor induction. This analysis focused on the 21 studies for which chromatin accessibility data were available in both conditions. An initial comparison of accessibility fold-changes between bHLH-bound and unbound regions revealed that CAT- and CAG-preferring factors—i.e., pro-differentiation bHLHs—exert detectable chromatin-remodeling activity across all cellular contexts tested (**Fig. S6A**). In contrast, factors within the CAC subclass showed no measurable chromatin-remodeling capacity (**Fig. S6A**).

In our previous work, we showed that strong central enrichment of a TF’s cognate motif at summits of highly dynamic ATAC-seq peaks is a hallmark of pioneer activity, indicating engagement and remodeling of previously closed chromatin (*64*). Here, for the subset of induction experiments with both pre- and post-induction ATAC-seq, we ranked peaks by fold-change, partitioned them into quartiles, and further subdivided each group according to the overlap with the corresponding TF’s ChIP-seq peaks. When assessing the centrality of Vierstra TF archetypes at the summits of fourth-quartile ATAC-seq peaks that overlap TF ChIP-seq peaks, E-box and E-box–like motifs consistently emerged among the most centrally enriched motifs across studies (**Fig. S6B**). E-boxes were also the most centrally enriched motifs in remodeled ATAC-seq peaks lacking TF binding, albeit with weaker enrichment (**Fig. S6C**). Notably, in the astrocyte-to-neuron conversion induced by NEUROG2, NFI motifs showed striking centrality in highly remodeled regions not bound by the factor (**Fig. S6C)**.

Next, we examined how the enrichment of E-box variants at ChIP-seq peak summits relative to flanking regions relates to chromatin remodeling. Using the same multivariate framework described above, we could determine that enrichments of each factor’s preferred E-box variants were positively associated with the extent of chromatin remodeling (**Fig. 5A**). However, this relationship largely disappeared after accounting for baseline pre-induction chromatin accessibility (**Fig. S7A**), suggesting that although specific E-box variants facilitate bHLH binding to closed chromatin, in general they do not determine the magnitude of the remodeling response.

**Fig. 5.**
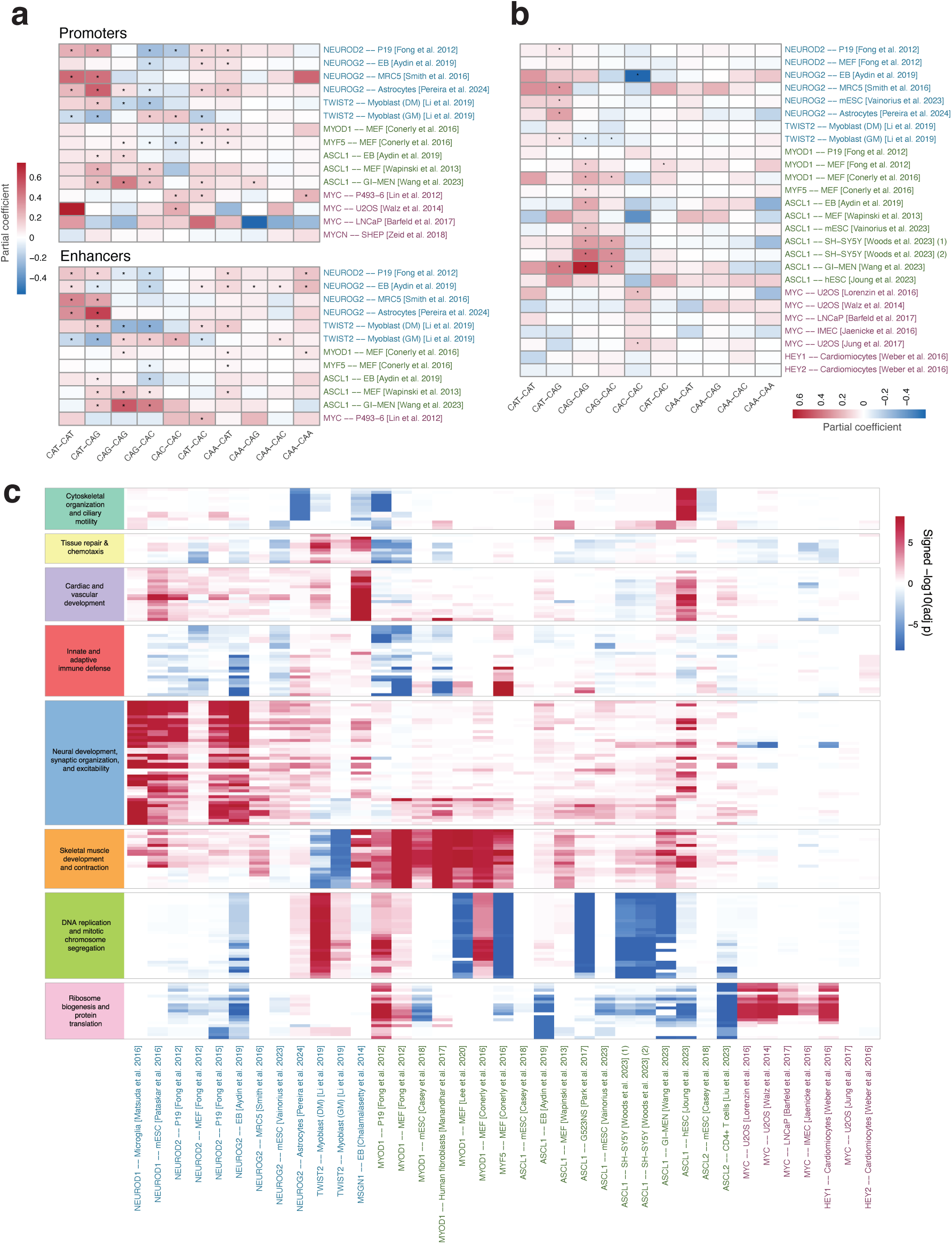
Molecular logic of the epigenomic and transcriptomic rewiring of the induced cells. **A**) Heatmap displaying the partial coefficients derived from multivariate linear models predicting chromatin accessibility fold-change based on the enrichment of specific E-box variants. The analysis is stratified by genomic location, discriminating peaks located within proximal promoters (<+-2kb from TSS) and distal enhancers (>+-4kb from TSS). Asterisks denote statistical significance after Benjamini-Hochberg correction (q < 0.05). **B**) Multivariate analysis of gene expression log2-fold changes relative to E-box enrichment. To ensure robust peak-to-gene association by proximity, the analysis was restricted to motifs located within proximal promoters (< +-2kb from the TSS). **C**) -log10 adjusted p values of gene ontologies, signed by the value of the NES. Only ontologies within the major clusters derived from figure S7 are included.

### Relationship between motif architecture and transcriptional activation

We next assessed whether motif architecture influences transcriptional output by analyzing post-versus pre-induction RNA-seq data and computing gene expression fold changes for genes including ChIP-seq peaks in their promoters (**Fig. 5B**). In contrast to chromatin remodeling, enrichment of E-box motifs in peaks remained positively associated with transcriptional activation even after adjusting for baseline chromatin accessibility (**Fig. S7B**). This ‘mass effect’ of E-boxes on gene transactivation was consistent across bHLH subclasses and was driven by their respective high-affinity E-box types. Proneural factors NEUROD2 and NEUROG2 drove transactivation using their high-affinity CAT-CAT and CAT-CAG E-boxes, whereas minor transactivation effects are detected for the CAG-CAG motif, which is broadly bound by these factors, but with a lower physical strength (*50*, *64*). Accordingly, the highest transactivation effect among MYC-bound E-box variants was observed for the high-affinity CAC-CAC, and not at the enhancer-enriched suboptimal CAT-CAC motifs. In contrast, no such association was detected for HEY factors, in line with their established function as transcriptional repressors (*74*). In summary, these results support a model in which local E-box density at binding sites enhances transcriptional activation independently of the extent of chromatin remodeling.

### Cross-study integration reveals TF-specific transcriptional programs

Finally, we integrated the transcriptional profiles across all induction studies. Clustering based on gene-expression fold changes revealed that experiments grouped primarily by the induced transcription factor rather than starting cell type or study origin (**Fig. S7C)**, underscoring the consistency with which each bHLH factor imposes its regulatory program. A particularly striking observation was that TWIST2 overexpression in myoblasts generated a transcriptional program inversely correlated with that of most CAT and CAG factors, including that of MYOD1, mirroring TWIST2’s established role in antagonizing myogenic differentiation and promoting a more undifferentiated cellular state.

Gene ontology analyses of the expression fold-change data further highlighted factor-specific transcriptional outcomes across cell types and studies. As expected, the strongest neurogenic signatures emerged following induction of the classical excitatory proneural factors NEUROD1, NEUROD2, and NEUROG2, whereas MYOD1 and MYF5 drove transcriptional programs associated with myogenesis. These proneural and myogenic factors shared transcriptional targets related to the regulation of neuromuscular functions and action potential (**Fig. S8**). Notably, ASCL1—though classically defined as a proneural factor— elicited a transcriptional program occupying an intermediate position between neurogenic and myogenic identities. Beyond developmental roles, ASCL1 exerts a distinct regulatory effect in cancer cells, where its overexpression consistently leads to the repression of proliferation-associated gene sets. Conversely, MYC overexpression leaves proliferation-associated programs largely unchanged, while selectively upregulating pathways related to protein translation and ribosome biogenesis, (**Fig. 5C, Fig. S8)**, suggesting that at oncogenic expression levels MYC primarily enhances biosynthetic capacity rather than cell-cycle progression. Together, these analyses reveal both specialization within bHLH subclasses and convergence across subclasses in the transcriptional programs they drive, indicating that functional outcomes cannot be explained solely by E-box preference but instead arise from factor-specific activity operating across diverse cellular contexts.

## DISCUSSION

### A unified, multi-omic framework for decoding bHLH regulatory mechanisms

Despite decades of work on bHLH transcription factors, the principles governing how different family members select genomic targets across chromatin contexts have remained unclear. Here, by systematically analyzing a comprehensive collection of bHLH induction ChIP-seq experiments, we define a DNA-binding code that explains this selectivity across the bHLH family. To gain a deeper insight into the molecular basis of bHLH activity, we integrated all bHLH binding profiles with multi-layered omic data, including chromatin accessibility, nucleosome positioning, CpG methylation, and transcriptomics, and assessed how the sequence determinants we have established—with particular emphasis on E-box variants defined by their central dinucleotide—vary across different molecular layers. These analyses were further complemented with HT-SELEX/methyl-SELEX data and structural modeling, allowing us to validate biochemical motif preferences and investigate the physical constraints of chromatin engagement. This multi-omic, cross-study comparative analysis uncovered new principles of bHLH binding logic, reconcile divergent observations across individual reports, and allowed findings to be generalized across bHLH subfamilies.

### Divergent strategies for engaging closed chromatin across CAT- and CAG-preferring bHLH factors

Several CAT- and CAG-preferring bHLH transcription factors—including MYOD1, ASCL1, NEUROD1, and NEUROG2—have previously been described as pioneer factors based on their ability to bind inaccessible chromatin and initiate lineage reprogramming (*4*, *5*, *9*, *46*, *64*, *90*, *91*). However, whether pioneer activity reflects intrinsic properties of individual factors or shared mechanisms within bHLH subclasses has remained unclear.

Building on our previous observations that NEUROD2 and NEUROG2 preferentially engage closed chromatin through CAT–CAT and CAT–CAG E-box variants (*64*), the present cross-study analysis supports a unifying model in which this mode of chromatin engagement extends broadly across CAT-preferring bHLH factors. Rather than supporting factor-specific distinctions in pioneer capacity, our cross-study comparison indicates that NEUROD1, NEUROD2, and NEUROG2 behave comparably across diverse cellular contexts. These findings challenge earlier models proposing differential pioneer activity within the proneural subgroup (*50*), and instead suggest that closed-chromatin engagement is a shared property of CAT-preferring bHLHs, manifested through common sequence features and chromatin remodeling signatures.

We further establish that CAG-preferring factors consistently use CAG–CAC and/or CAG–CAG E-box variants to engage closed chromatin. Although a previous study reported a differential enrichment of CAG–CAC and CAG–CAG motifs in closed versus open chromatin, respectively (*9*), our cross-study analysis indicates that this relationship is not universal and varies across experimental contexts.

An exception to the general behavior of CAT- and CAG-preferring bHLH factors is the TWIST family. We extend previous observations of 5-bp E-box spacing (*25*, *92*), showing that this configuration is uniquely associated with TWIST factors and is selectively enriched when they engage closed chromatin. Consistent with this distinctive binding architecture, and in line with previous findings that TWIST2 antagonizes MYOD1 during myogenesis (*25*), we found that the transcriptional program induced by TWIST2 is opposite to those driven by other CAG and CAT-preferring factors. Together, our integrative analyses provide an explanation of why, unlike their phylogenetically close CAT- and CAG-preferring factors that broadly promote cell differentiation, TWIST factors instead exert their well-established roles in cell dedifferentiation and cancer progression (*25*, *26*, *56–58*, *93*, *94*).

### A reevaluation of the model of bHLH nucleosome interaction

A widely cited model has linked bHLH pioneer capacity to the length of their basic α-helix, proposing that shorter helices better avoid steric clashes with the nucleosome and thus enable binding to nucleosome-engaged DNA (*95–98*). This model further postulates that degeneracy at the central dinucleotide of the E-box arises as a direct consequence of α-helix length; bHLHs with longer α-helices establish full sequence-specific contacts with the E-box, which is reflected in strong conservation of the central dinucleotide. Conversely, bHLHs with shorter α-helices are thought to reduce or lose contacts at this position, leading to increased degeneracy at the intermediate bases of their motifs. Our results do not support an association between α-helix length, E-box degeneracy, and pioneer activity. Across CAT- and CAG-preferring bHLHs, we observed comparable ability to access low-accessibility chromatin despite substantial intra-group variation in helix length. With respect to the degeneracy of the central dinucleotides linked to pioneer activity, our results point to the opposite direction as of the aforementioned model, suggesting that the bHLH factors preferentially bind to their preferred high-affinity E-box variants when targeting less accessible chromatin. Furthermore, nucleosome chemical-mapping data demonstrates that, despite differences in helix length, all bHLH factors bind preferentially at nucleosome flanks rather than at the dyad center, in line with recent studies (*89*, *99*). Together, these findings argue against helix length or E-box degeneracy as primary determinants of nucleosome engagement for bHLH factors.

### An E-box–affinity–dependent model of chromatin remodeling and transcription

Instead, we propose a model in which the ability to act as pioneer factors—accessing and remodeling closed chromatin– is a shared property of CAT and CAG-preferring bHLH factors and is largely absent from the evaluated CAC-preferring bHLH subclass. We further suggest that pioneering activity is E-box-variant dependent. CAG-preferring bHLH factors preferentially engage closed chromatin through CAG-CAG or CAG-CAC E-boxes. CAT-preferring factors show a marked enrichment for CAT-CAT/CAT-CAG in inaccessible regions, whereas their broadly bound CAG-CAG motif is largely invariant with respect to chromatin accessibility.

Remarkably, a similar E-box preference is evident at the level of transcriptional activation of nearby genes. CAG-preferring factors preferentially transactivate through CAG–CAG and CAG–CAC E-boxes, whereas CAT-preferring factors act predominantly through CAT–CAT and CAT–CAG motifs. In vitro assays have shown that CAT–CAT (*64*) and/or CAT–CAG (*100*) motifs represent the highest-affinity binding sites for CAT-preferring factors; and that CAG–CAG/CAG–CAC motifs are optimal for CAG-preferring bHLHs (*76*, *101*, *102*). Because higher DNA-binding affinity has been shown to enhance both competition with nucleosomes (*103*) and recruitment of chromatin remodelers and transcriptional coactivators (*104*, *105*), we propose an E-box–affinity–dependent model of epigenomic and transcriptomic rewiring. In this framework, differential residence times at specific E-box variants enable distinct degrees of chromatin remodeling and transcriptional activation, without requiring a strict coupling between the two processes.

Consistent with this model, we observe increased motif density in closed chromatin, in line with the known effect of motif clustering on binding affinity. Likewise, TWIST binding to inaccessible regions is selectively enriched for 5-bp–spaced E-boxes, a configuration known to enhance DNA binding through tetramerization.

Across all datasets, E-box and E-box–like motifs are the dominant sequence centrally enriched at both ChIP-seq summits and remodeled ATAC-seq peaks, indicating that bHLH factors primarily rely on intrinsic DNA-binding specificity rather than obligate cofactor motifs to engage closed chromatin. This stands in contrast to interpretations from some previous studies, which proposed that bHLH factors depend on non-bHLH cofactor motifs to achieve binding specificity (*10*, *50*, *106–108*) or to access closed chromatin (*95*, *109*, *110*). Together, these observations indicate that increased DNA binding strength—achieved either through high-affinity E-box-variant binding, motif clustering or specific spacing architectures—facilitates bHLH engagement with closed chromatin.

### A methylation-driven model explaining MYC enhancer invasion

By integrating chromatin, methylation, and hexanucleotide-resolution binding information across all E-box variants, we delineate a mechanistic model explaining why CAC bHLH factors (MYC, MYCN, MITF) invade enhancers when overexpressed in multiple cancer types (*17*, *63*, *111–116*). Under physiological expression levels, MYC preferentially binds its optimal CAC–CAC E-boxes predominantly located in accessible promoters. However, when overexpressed, we show that MYC can invade enhancers with high CpG methylation levels, shifting its preference from CAC-CAC to CAT-CAC E-boxes. Indeed, it has been noted a preferential binding of the CAT-CAC E-box when invading enhancers, in contrast to the endogenously bound CAC-CAC in promoters (*17*, *63*, *111*, *113*), which is consistent with earlier evidence that when overexpressed—a hallmark of many tumor types—MYC can engage lower-affinity E-box variants beyond CAC-CAC (*78*, *111*, *117*). By interrogating *in vitro* methyl-SELEX data, we demonstrate that this shift in E-box usage corresponds to a change in MYC’s intrinsic DNA-binding preferences: when the sequences are methylated, MYC binding to the CpG-containing CAC-CAC (CACGTG) E-box is abolished, and it instead binds DNA through the suboptimal CAT-CAC E-box. Together, our observations point to a model where at oncogenic expression levels, MYC expands its binding landscape by targeting CAT-CAC E-boxes within highly methylated enhancer regions where CAC-CAC sites are otherwise “blocked.”

Interestingly, CAT–CAC motifs are also frequently enriched in TWIST-binding regions, and both TWIST and MYC/MYCN/MITF can activate pro-oncogenic transcriptional programs. This convergence suggests that the ability to use CAT-containing motifs in methylated regulatory regions may represent a shared mechanism for accessing oncogenic enhancers. In line with this, TWIST1 ChIP-seq peaks have been found colocalizing with MYCN in MYCN-invaded enhancers in neuroblastoma cell lines (*63*).

Similarly, we find that HEY factors also undergo a previously unrecognized, methylation-dependent shift in DNA binding, whereby their canonical preference for CAC–CAC E-boxes becomes redirected toward CAT-containing motifs under high DNA methylation. Notably, HEY proteins are strikingly elevated across multiple aggressive tumor types and their aberrant expression is associated with poor prognosis and adverse overall survival (*118*). Together with the behavior of MYC and TWIST, these observations suggest potential cross-talk among MYC, TWIST, and HEY family members at low-affinity CAT–CAC motifs within oncogenic enhancers.

### Interpreting bHLH activity under induced conditions

We have focused our study on single-factor induction assays, which has provided a controlled framework to isolate the intrinsic activity of individual transcription factors. By uncoupling TF action from higher-order regulatory interactions, this approach enables direct assessment of DNA-binding preferences, i.e. chromatin engagement modes, and motif features —such as E-box type, density, and spacing—that are inherent to each bHLH factor or subclass. The persistent emergence of these subclass-specific binding architectures across diverse datasets—transcending both technical noise and cellular context—allows us to confidently delineate a conserved regulatory grammar for the bHLH family.

Admittedly, while these assays reveal intrinsic regulatory logic, they inevitably bypass the full complexity of physiological TF networks and the multifaceted environmental cues that modulate transcriptional responses *in vivo*. Moreover, increased TF dosage can artificially promote binding to otherwise inaccessible sites, as exemplified by TWIST1 overexpression (*119*). In the context of oncogenic bHLH factors such as TWIST or MYC family members, overexpression is nevertheless particularly informative, as it mirrors pathological scenarios in which elevated TF levels drive enhancer invasion and regulatory rewiring. By contrast, for proneural factors, forced induction can perturb stoichiometric relationships with endogenous partners—especially E-proteins—thereby redirecting binding from heterodimer-preferred to homodimer-preferred E-box variants (*64*, *120*). Differences in TF concentration therefore emerge as critical determinants of the bHLH binding landscape and, ultimately, of lineage commitment. Importantly, analyses based on endogenous in vivo expression of NEUROD2 and NEUROG2 revealed comparable—and in some cases stronger—associations between CAT–CAT and CAT–CAG E-box enrichment and chromatin accessibility, supporting the robustness of these relationships beyond induced conditions (*64*).

Intriguingly, the ability of many bHLH factors to remodel chromatin does not fully explain their cell type–specific activity, as even established pioneer factors can trigger vastly different tissue identities depending on the initial cellular environment (*10*, *121*). For example, NEUROD1 can drive either neuronal or pancreatic differentiation depending on the induced cell type (*122*), and PTF1A can similarly specify both lineages in a manner dependent on the presence of the cofactor RBPJ (*123*). These observations suggest that additional molecular and epigenetic, context-dependent mechanisms modulate how bHLH factors engage DNA and initiate lineage-specific transcriptional programs.

### Conclusion and future directions

Taken together, this work provides a unified framework for understanding how bHLH transcription factors interpret the E-box code across variable chromatin environments and demonstrates that much of their capacity to engage closed chromatin is encoded directly in the sequence architecture of the motifs they recognize. By integrating genomic, epigenomic, and structural data, we uncover both shared principles and subfamily-specific mechanisms, offering a coherent explanation for the functional diversity of bHLH factors and a foundation for interpreting new in vivo datasets. Importantly, in vivo delivery of bHLH factors via viral vectors has demonstrated reprogramming capacity and therapeutic potential in multiple mouse disease models, such as diabetes (*124–126*), obesity (*127*), Parkinson’s disease (*128*, *129*), Alzheimer’s disease (*34*), and different types of cancer (*55*, *130–134*). As future in vivo induction experiments and high-resolution chromatin profiles become available, this framework will serve to predict, model, and potentially modulate bHLH–chromatin interactions in development, reprogramming, and disease.

## Methods

### ChIP-seq peaks QC and general motif specificity description

ChIP-seq peak sets were retrieved from the ChIP-Atlas database (https://chip-atlas.org/) (*135*) for all studies in which a single bHLH TF had been ectopically overexpressed in cultured cells (**Table S1**). When multiple inductions of the same TF were performed in the same cell type (biological replicates), each induction was treated as an independent experiment. When a ChIP-seq experiment (biological replicate) included more than one technical replicate, a high-confidence set of peaks was derived by retaining only those peaks detected in at least two replicates. The peaks intersecting with ENCODE blacklist genomic regions were excluded from all analyses. In cases where ChIP-seq data were available at multiple time points after TF induction, the dataset with the largest number of peaks was selected (**Table S3**).

To further refine the quality of ChIP-seq datasets, we assessed the positional enrichment of E-box motifs around peak summits. All peaks were resized to 400 bp centered on the summit, and E-boxes were identified within this window by simple pattern matching of the canonical CANNTG sequence. The centrality of motif positions relative to the summit was then quantified using a one-sided Kolmogorov–Smirnov (K–S) test, comparing the empirical distribution of distances between E-boxes and summits to a uniform null distribution spanning –199 to +199 bp. Under the null hypothesis, motif positions are uniformly distributed across the window, whereas the alternative hypothesis (“less”) tests for an overrepresentation near zero, indicating central enrichment. A low p-value (p < 0.05) therefore reflects a significant clustering of E-boxes around the summit. ChIP-seq experiments were ranked by K–S p-value (centrality), and the bottom 15 experiments showing the weakest enrichment were excluded from subsequent analyses.

For each set of ChIP-seq peaks, number of motifs per peak and spacing distance among E-boxes were computed using regular expression-detected E-boxes mapped within original coordinates of the peaks. Motif centrality was computed using peaks resized 400bp around the summit, as the inverse of the median distance of each E-box variant from the peak summit. To calculate the centrality of Vierstra motif archetypes (v1.0), BED files containing the genomic occurrences of each motif in the mm10 and hg38 assemblies were downloaded from (https://www.vierstra.org/resources/motif_clustering#downloads) and intersected with our 400bp-resized ChIP-seq peaks.

### Pre induction chromatin

For the pre-induction chromatin accessibility analyses, we obtained chromatin accessibility data matching the cell type of each induction experiment (**Table S4**). When available, ATAC-seq data was prioritized over histone ChIP-seq datasets. In the absence of ATAC-seq, H3K27ac ChIP-seq was selected preferentially over ChIP-seq data for other histone post-translational modifications. And whenever the same study that generated the TF ChIP-seq data also provided its own ATAC-seq dataset, that dataset was used.

All chromatin accessibility datasets were reprocessed using an in-house pipeline. FASTQ files were downloaded from the SRA using **fasterq-dump** (part of SRA Toolkit v3.1.1) with the corresponding accession numbers. Reads were aligned to the mm10 and hg38 reference genomes using **bwa-mem (v0.7.17) [Li, 2013]** with default parameters. Duplicate reads were removed with **samtools rmdup** (part of SAMtools v1.21) (*136*), and only uniquely mapped reads were retained using the following filter: `grep -v -e ‘XA:Z:’ -e ‘SA:Z:’`. Aligned BAM files were then converted to bedGraph format using the `bamCoverage` function from **deepTools (v3.5.5)** (*137*), with normalization set to RPKM (`--normalizeUsing RPKM`). To integrate chromatin accessibility with ChIP-seq data, the ChIP-seq peaks from each experiment were resized to 1,000 bp centered on the summit. The resulting bedGraph files were intersected with these resized peaks using the `bedtools map` function (**BEDTools v2.31.1**) (*138*) with the `-max` parameter, assigning to each ChIP-seq peak a pre-induction chromatin accessibility value corresponding to the maximum signal within the 1,000 bp window around the summit.

To quantify the influence of specific E-box variants on chromatin accessibility, we developed a computational framework to calculate a motif enrichment index for each of the ten E-box variants. For each ChIP-seq peak, motif occurrences were identified within the peak coordinates and compared against the local genomic background to account for regional sequence biases. This background was defined by extracting the immediate upstream and downstream flanking regions of each peak, the sum of the upstream and downstream background sequences being the same as the length of the peak. To stabilize the variance and account for peaks with zero counts, we applied a pseudocount of +1 to both the peak and flanking counts. The motif enrichment index was then calculated as the log-ratio of motif density within the peak relative to its flanking background: motif enrichment index= log[(count_peak+1) / (count_flanking+1)].

The relationship between motif enrichment and chromatin accessibility was assessed using a multivariate linear regression approach. This continuous model allowed for the simultaneous evaluation of all ten E-box variants as independent predictors. For each study, we fit a linear model where the log-transformed accessibility of the peak served as the dependent variable and the enrichment indices of the E-box variants were used as predictors.

Statistical significance was determined by calculating the p-values associated with the regression coefficients (Beta) for each motif. To account for multiple hypothesis testing across all studies and motifs, p-values were adjusted using the Benjamini-Hochberg (BH) procedure. Finally, to visualize the relative contribution of each E-box variant to chromatin accessibility across different experimental conditions, the partial coefficients were plotted in a heatmap. In these visualizations, significant associations (defined by a False Discovery Rate, q < 0.01) were highlighted with an asterisk.

### Methylation

Methylation data for the available cell types (Table S6) were obtained as preprocessed files containing both CpG genomic coordinates and their corresponding methylation percentages. In some cases, CpG methylation data were aligned to the mm9 or hg19 assemblies; these coordinates were converted to mm10 and hg38, respectively, using the **liftOver** tool (*139*). The methylation data were then intersected with the original ChIP-seq peak coordinates using the `bedtools map` function (**BEDTools**) (*138*) with the `-mean` parameter, to compute the average CpG methylation percentage within each peak.

### Methyl-selex

HT-SELEX and methyl-SELEX data were obtained from Yin *et al.* (2017) (*140*). After downloading the FASTQ files for each successive SELEX round, we used regular expression matching to identify E-box variants and calculated the proportion of sequences containing each E-box type per round.

### Nucleosome maps

To investigate the spatial relationship between bHLH binding sites and nucleosome organization, we used high-resolution chemical nucleosome mapping data from mESC, provided by Voong et al., as BED files of the dyad positions (GEO: GSM2183909) (*86*). This approach provides base-pair–resolved nucleosome dyad positions with reduced sequence bias compared to MNase-based methods, particularly in A/T-rich genomic regions.

Dyad positions originally mapped to the mm9 genome assembly were lifted over to mm10 using the UCSC liftOver tool. Nucleosomal regions were defined by extending dyad coordinates ±100 bp. ChIP-seq peak summits for bHLH factors, as well as the pioneer factors OCT4 and SOX2 (SRA accessions SRX236476 and SRX236477, respectively), were intersected with the +-100bp-extended dyads, and base-pair distances relative to the dyad were computed using the valr package (*141*). For each dataset, summit frequencies were quantified at 1-bp resolution and min–max normalized to enable comparison across experiments. Profiles were smoothed using a Gaussian kernel (window = 11 bp, SD = 3).

### ModCRE

Structural models of bHLH dimers bound to nucleosomal DNA were generated using the ModCRE package (*142*), specifically the complexbuilder module from the ModCRElib library (*142*). The workflow consisted of three main stages.

First, a series of 146-bp nucleosomal DNA sequences was constructed in which the canonical bHLH binding motif (CATATG) was systematically positioned at every location from nucleotide 1 to 140. The structure of each sequence was then modeled in a nucleosomal conformation using X3DNA, with the crystal structure of a canonical nucleosome (PDB ID: 6FQ5 (*143*)), including histone proteins, serving as the structural template.

Second, structural models for all bHLH dimers—both homo- and heterodimeric combinations—were generated with AlphaFold3 (*144*) using DNA sequences containing the CATATG motif. The resulting models were trimmed to remove unstructured terminal regions, retaining only well-defined and contiguous secondary-structure elements corresponding to the DNA-binding domains of the transcription factors.

Third, each bHLH dimer was positioned onto each nucleosomal DNA conformation using the complexbuilder module. Clash detection was performed between backbone atoms of the nucleosome (histones plus DNA, excluding the binding motif) and backbone atoms of the TF DNA-binding domains, using distance thresholds ranging from 0.5 Å to 6.0 Å in 0.5-Å increments. A maximum of two steric clashes per model was permitted; configurations exceeding this limit were excluded from further analysis.

Transcription-factor occupancy along the nucleosomal DNA was quantified by sliding the CATATG window from positions 1 to 146 and recording, at each position, whether the nucleotide was in contact with the TF. Contacts were defined using the standard ModCRE criterion of <4 Å between any atom of the TF and any atom of the corresponding nucleotide. Because of variations in the length and orientation of the bHLH binding helices, some dimers exhibited strong periodicity, binding preferentially at positions where the helices could access solvent-exposed DNA regions. Under the more permissive 6-Å contact threshold, most dimers displayed a characteristic repeating pattern corresponding to outward-facing DNA grooves that recur approximately every 6 bp along the nucleosome. Finally, TF occupancy at the 4-Å threshold was quantified for all models, enabling direct comparison of nucleosomal binding patterns across different homo- and heterodimeric bHLH combinations All structural models were visualized using Chimera (*145*).

### Chromatin remodeling

For all studies with chromatin accessibility data available both before and after induction, peak calling was performed to identify accessible regions using the BAM files generated as described above. Peak calling was carried out with **MACS2 callpeak (v2.2.9.1)** (*146*), using the `--narrow` parameter for ATAC-seq data and the `--broad` parameter for histone PTM ChIP-seq data. Pre- and post-induction peak sets were merged into a single set, and bedGraph files containing pre- and post-induction coverage were mapped onto the merged peak set using the `bedtools map` function (**BEDTools**) with the `-max` parameter. This allowed the computation of fold changes in chromatin accessibility between pre- and post-induction conditions for each peak. The resulting accessibility peaks were then intersected with the corresponding transcription factor ChIP-seq peaks.

To assess E-box influence on chromatin accessibility variation within TF-bound regions, we modified the mapping approach to account for peaks missed by the initial peak caller. ChIP-seq peaks were resized to 1,000 bp centered on the summit, and pre- and post-induction accessibility signals were mapped directly onto these windows using the maximum value. This ensured that fold changes were computed for all TF-bound regions, even those that did not intersect with the previously generated chromatin accessibility peak sets. Then, the log2 fold change was computed for the post-vs pre-induction chromatin accessibility. Peaks were stratified by genomic location into promoters, defined as regions within +-2kb of a Transcription Start Site (TSS), and distal enhancers, located more than 4kb away from a TSS. Only datasets containing a minimum of 1,000 peaks were included in the downstream statistical modeling to ensure sufficient power and robust estimation of effect sizes.

We employed multivariate linear regression to model the relationship between the enrichment of the E-box variants and the degree of chromatin remodeling. To isolate the intrinsic potency of specific motifs from the confounding influence of the pre-existing chromatin state, we implemented two parallel modeling strategies for both promoters and enhancers. The basic model evaluated accessibility changes as a function of the ten motif enrichment indices, while the adjusted model included initial (pre-induction) log-transformed accessibility as a covariate. Regression coefficients (Beta) were estimated for each motif, and p-values were adjusted for multiple testing using the Benjamini-Hochberg procedure to maintain a False Discovery Rate (FDR) below 0.01.

### Transcriptomic Analysis

Raw RNA-seq FASTQ files were retrieved from GEO and SRA archives. RNA-seq quality control and preprocessing were performed using fastp (*147*) and FASTQC. Files with very low read counts (<1million reads) were excluded from the analysis. Preprocessed reads were aligned to the reference genome using STAR (*148*). GENCODE 47 (GRCh38.p14, human) and GENCODE M36 (GRCm39, mouse) annotation genomes were employed. Mapped reads were quantified using FeatureCounts (*149*). For single-end reads, default parameters were applied. For paired-end reads, fragments were counted instead of individual reads, and only fragments where both ends were mapped were included. This approach was implemented to improve comparability between the two data types. Differential expression analysis for RNA-seq data was performed using DESeq2 (*150*) and contrasts were calculated between bHLH transcription factor-induced and uninduced samples. In most cases, comparisons were made within the same replicate before and after induction, treating the analysis as a paired design. For microarray data, differential expression analysis was conducted using Limma (*151*). Intensities of probes corresponding to the same genes were averaged.

To integrate transcriptional responses across different cell types and species, we first identified high-confidence orthologs using the biomaRt R package to map Ensembl gene IDs and symbols between *Homo sapiens* and *Mus musculus*. A joint expression matrix was generated by merging the datasets based on these shared orthologs, where rows represented orthologous genes and columns represented the individual studies. Pearson correlation coefficients were then calculated for all pairs of studies using a pairwise complete observation approach. The resulting correlation matrix was visualized as a heatmap using the pheatmap package, employing a divergent color scale from blue (negative correlation) to red (positive correlation) and organized via hierarchical clustering to identify clusters of transcription factors and experimental conditions that exhibit highly coordinated gene expression responses.

To characterize the functional transcriptional responses across datasets, Gene Set Enrichment Analysis (GSEA) was performed using the clusterProfiler package (*152*). Genes from each study were ranked by their log2 fold-change and assessed against the Gene Ontology: Biological Process (GO-BP) database, utilizing org.Hs.eg.db for human and org.Mm.eg.db for mouse studies. The results were integrated into a balloon plot representing a master list that included the top 10 most significant ontologies from each study. In this visualization, the size of each dot is proportional to the statistical significance, while the color represents the Normalized Enrichment Score (NES) using a divergent scale to distinguish between activated (red) and suppressed (blue) pathways. Finally, ontologies were organized via hierarchical clustering based on their NES profiles and manually assigned to functional categories to highlight shared regulatory programs across transcription factors and cell types.

To investigate the association between E-box variant enrichment and expression activation of genes, the same strategy as in the chromatin remodeling was followed, substituting chromatin accessibility fold change with expression fold change. To gain confidence, expression fold change was assigned only to the ChIP-seq peaks within a distance of +-2kb from the TSS of the gene.

## Acknowledgements

**Funding:** Agencia Estatal de Investigación /AEI/10.13039/501100011033 [PID2020-113203RB-I00 to B.O., PID2022-140137NB-I00 to G.S.]; Instituto de Salud Carlos III [CP20/00064], with co-financing by European Funds for the Miguel Servet Contract; Fundació la Marató de TV3; Unidad de Excelencia María de Maeztu [CEX2018-000792-M]; MCIN [PRE2020-093064 to X.d.M.]. Funding for open access charge: Instituto de Salud Carlos III [CP20/00064], with co-financing by European Funds for the Miguel Servet Contract. This work has been carried out within the framework of the PhD program in Genetics at the Autonomous University of Barcelona. **Authoŕs contributions**: X.d.M., JMM, SB, AN and JI performed the bioinformatic analyses for this study. X.d.M. and G.S. conceived the study and wrote the manuscript, with contributions of B.O. B.O. performed the ModCRE analyses. G.S. supervised the study. All authors read and approved the manuscript. **Data and materials availability:** All data needed to evaluate the conclusions in the paper are present in the paper and/or the Supplementary Materials. All code to reproduce the analyses presented in the paper is available at: https://github.com/SantpereLab/bHLH_Induction. This work has been carried out within the framework of the PhD program in Genetics at the Autonomous University of Barcelona.

---

**Table S1. Summary of *in vitro* bHLH induction studies found in the literature.** This table shows all the experiments found in the literature where expression of a single bHLH transcription factor was artificially induced, in *in vitro* cell cultures. The biological effect of the ectopic overexpression is shown as it was described in the referenced paper.

**Table S2. Summary of bHLH induction datasets included in this study.** Detailed description of all ChIP-seq datasets derived from bHLH induction experiments considered in our analysis, where each row represents each experiment. Only studies in which (i) the ChIP-seq was performed following the induction of a single bHLH transcription factor and (ii) no additional experimental perturbations were introduced were included.

**Table S3. Peak counts across ChIP-seq replicates.** Total peaks identified per for all bHLH datasets sourced from ChIP-Atlas. High-confidence peaks were defined by their presence in at least two replicates, excluding any regions overlapping ENCODE blacklisted sites. Each row of the table represents an individual ChIP-seq experiment; for multi-timepoint studies, each row corresponds to a specific timepoint. To capture the period of highest TF activity, only the timepoint with the highest number of identified peaks was retained for downstream analysis.

**Table S4. Chromatin accessibility data used for each cell line.** Original study citations and GEO accession numbers for each sample used in the pre-induction chromatin accessibility analysis of each cell line. For certain cell lines, multiple entries are listed; this occurs when more than one induction study provides pre-induction accessibility data for that cell line. Entries highlighted in red denote data sourced from studies outside of the primary bHLH induction datasets, where accessibility data from matching cell lines were incorporated.

**Table S5. Literature review of E-box motif analyses within the re-analyzed datasets.** For each ChIP-seq experiment included in this study, we assessed the depth of the motif analysis performed in the original publication. We categorized whether the authors analyzed E-box motifs using general Position Weight Matrices (PWMs) or if they stratified them by all central dinucleotide combinations. The third column specifies whether the motif analysis involved a comparison between different subsets of ChIP-seq peaks. Highlighted in red are the three studies that both analyzed the central dinucleotide of the E-boxes and compared motifs between pre-induction open versus closed chromatin.

**Table S6. Summary CpG methylation data sources.** Accession numbers of the CpG methylation data we found in the literature that matched the cell lines under study.

**Fig. S1.**
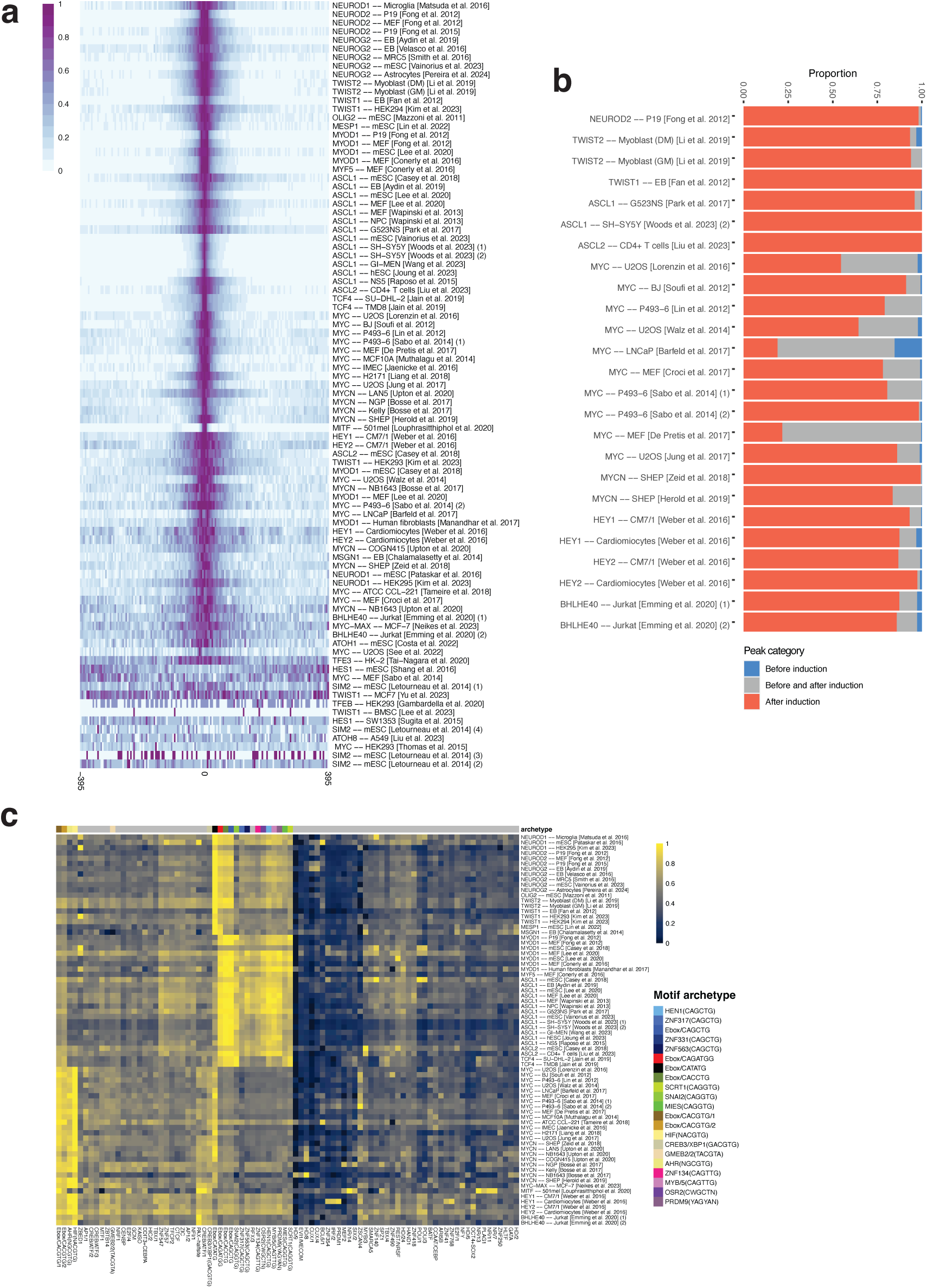
Preliminary characterization of motif enrichment and genomic distribution of ChIP-seq peaks. **A)** Heatmap displaying the density of canonical CANNTG E-box motifs binned at 5 bp resolution relative to ChIP-seq summits. Motif counts are min-max scaled within each experiment (row) to facilitate cross-study comparison. Experiments are ordered from the top to the bottom by motif centrality, determined by a Kolmogorov-Smirnov (KS) test comparing observed distributions against a theoretical uniform distribution. **B)** Stacked barplot illustrating the intersection of pre-induction and post-induction ChIP-seq peaks. Peaks are categorized as pre-induction specific (blue), persistent/shared (gray), or post-induction specific *de novo* binding (red). **C)** Heatmap showing the row-scaled centrality of Vierstra motif archetypes within +-200bp of peak summits. The analysis is restricted to a master list of motif archetypes derived from merging the top 50 most enriched archetypes in each study. The top annotation bar distinguishes canonical E-boxes and E-box-like motifs (colored) from non-E-box motifs (gray), based on manual curation. Values are row-normalized to emphasize study-specific preferences. **D)** Stacked barplot illustrating the proportion of peaks categorized as promoters, proximal enhancers, or distal enhancers. Comparisons are shown for pre-induction and post-induction peaks to highlight global shifts in the regulatory landscape

**Fig. S2.**
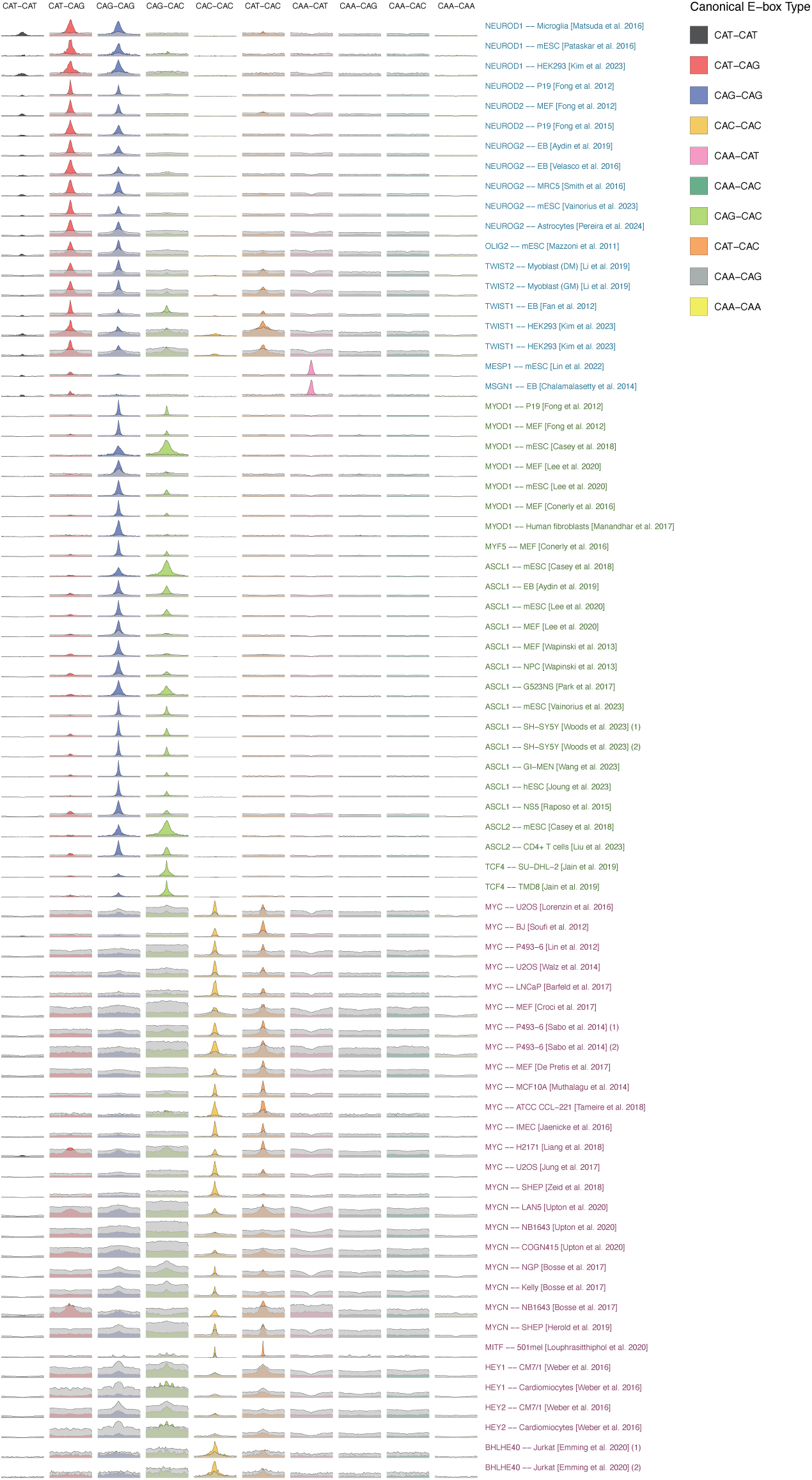
Detailed characterization of canonical and non-canonical E-box variant enrichment. **A)** Distribution of canonical and non-canonical E-boxes around the summits of ChIP-seq peaks, resolved by E-box variant. The distributions show frequencies of motifs per peak in a window of +-400bp around the summits, being these frequencies min-max normalized globally in each study. This allows for an intra-study comparison of the enrichment of different E-box variants, resolved by both their central NN dinucleotide and the presence of degeneracy at the canonical CA/TG positions.

**Fig. S3.**
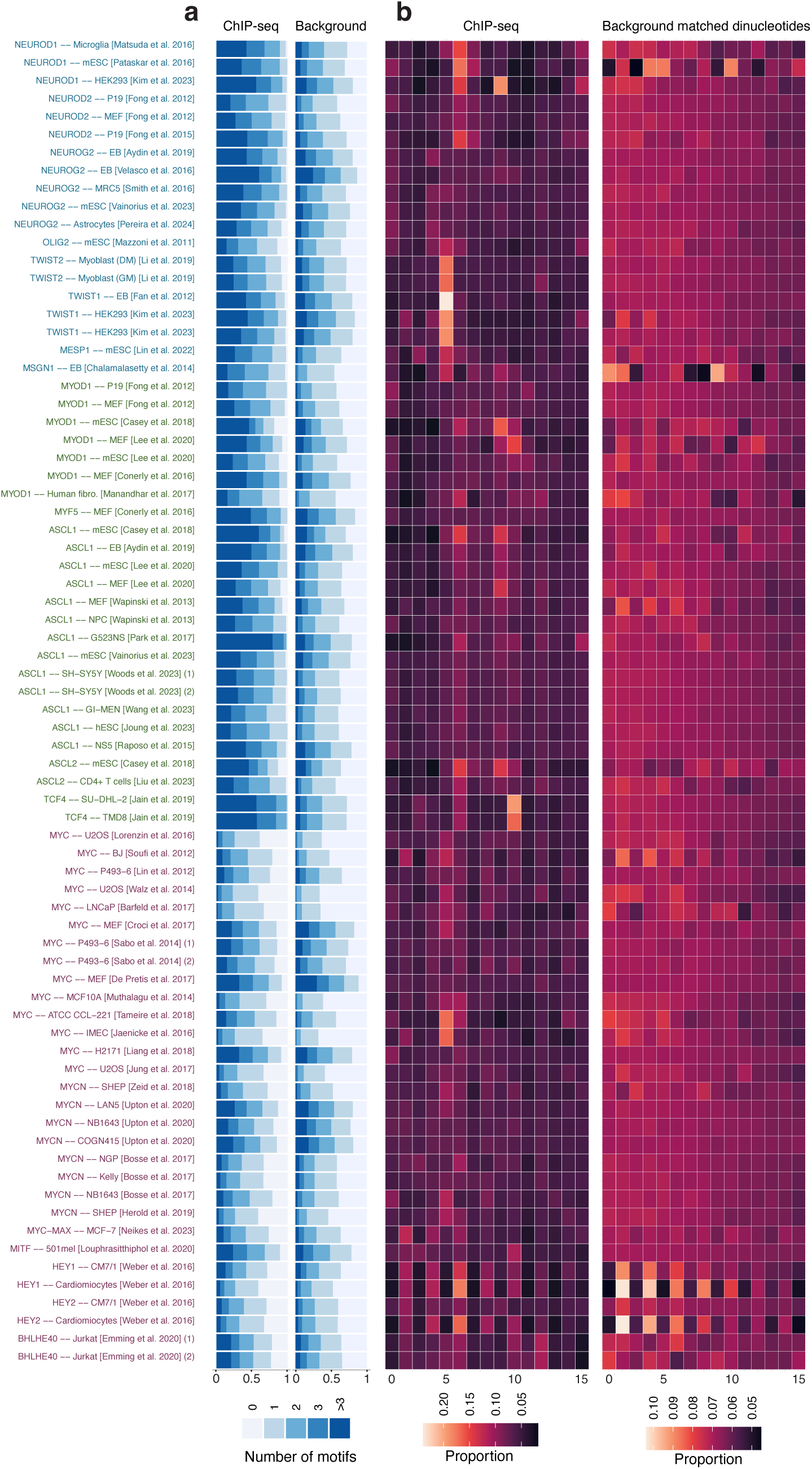
Assessing bHLH binding site architecture through background sequence comparison. **A)** Proportions of ChIP-seq peaks containing varying numbers of canonical E-boxes compared to flanking background sequences. Background regions were defined by extending half the peak length both upstream and downstream of each peak. **B)** Proportion of Xbp-spaced tandem canonical E-box motifs, in the ChIP-seq peaks (left) and in dinucleotide content-matched artificially generated background sequences (right).

**Fig. S4.**
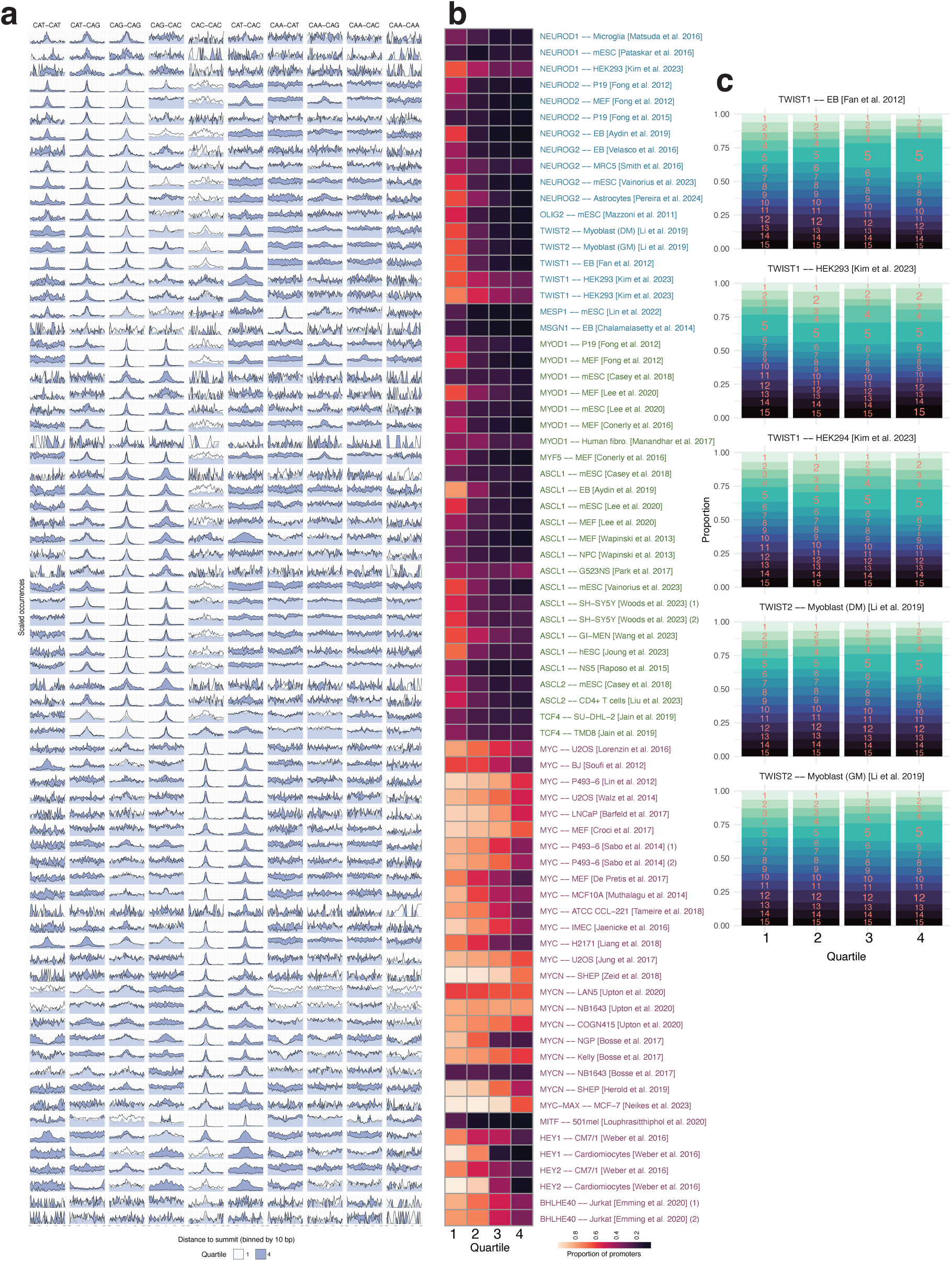
Inspection of DNA binding logic of bHLH factors in the pre-induction binding landscape. **A)** Distribution of motifs around summits of peaks, stratified by binding to open (Q1) and closed (Q4) chromatin. Distributions are scaled relative to the maximum of each study / E-box variant, to allow for intra-motif comparisons of Q1 vs Q4. **B)** Heatmap depicting the proportion of peaks that fall in promoter regions within each accessibility quartile in each study. **C)** Stacked barplot depicting the proportion of spacings between E-boxes in the accessibility quartiles of the TWIST1/TWIST2 induction experiments.

**Fig. S5.**
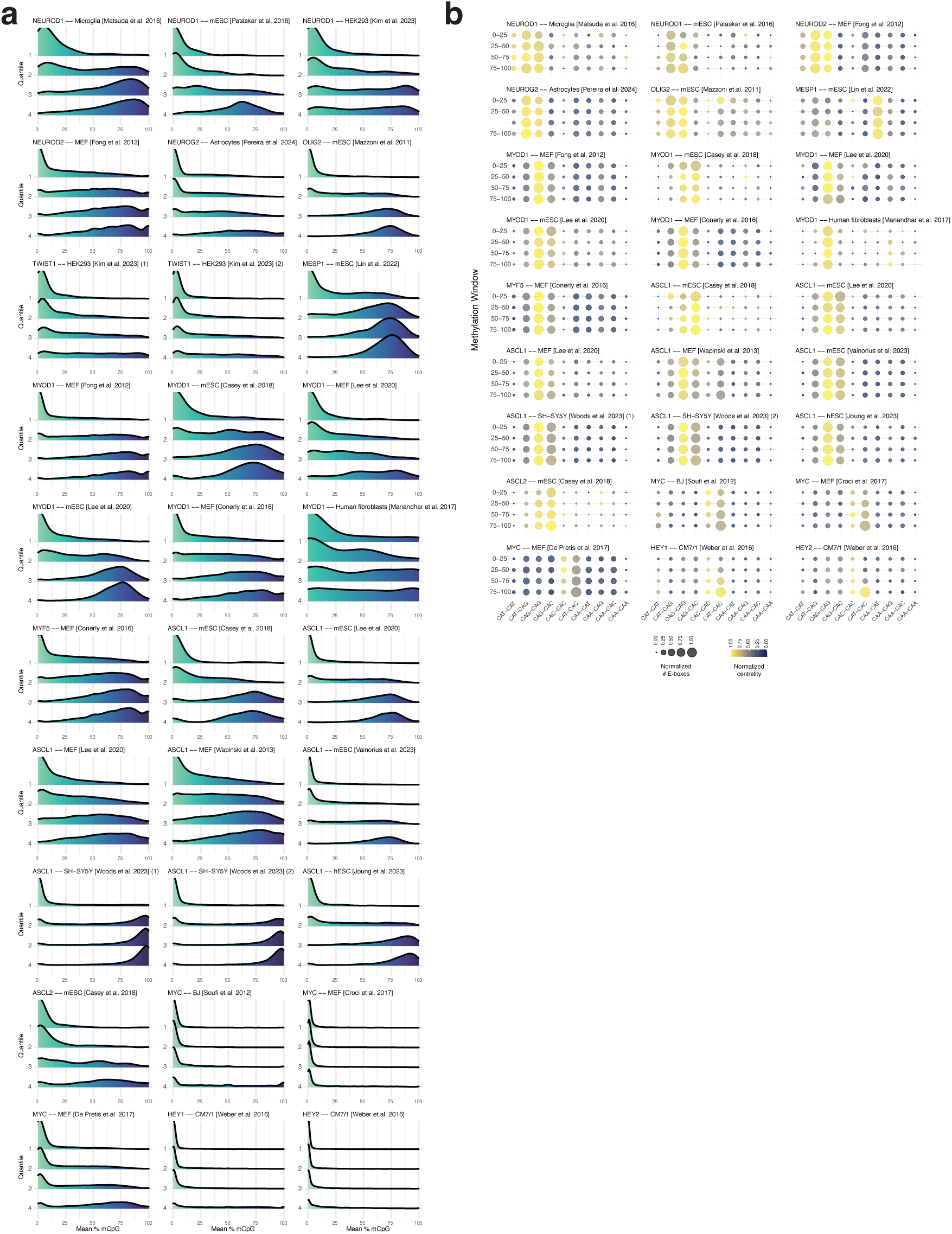
Global dissection of genomic targeting across differentially methylated regions. **A)** Chromatin accessibility quartile-resolved density plots, where the X axes illustrate the mean percentage of CpG methylation within ChIP-seq peaks. **B)** Balloon plots showing centrality and number of motifs per peak for each E-box variant, restricting the analysis to the Q4 of chromatin accessibility (less accessible), and further stratifying the peaks in %CpG methylation bins. Balloon size and color are min-max scaled within each balloon plot (each study), to allow comparisons between motifs within each study.

**Fig. S6.**
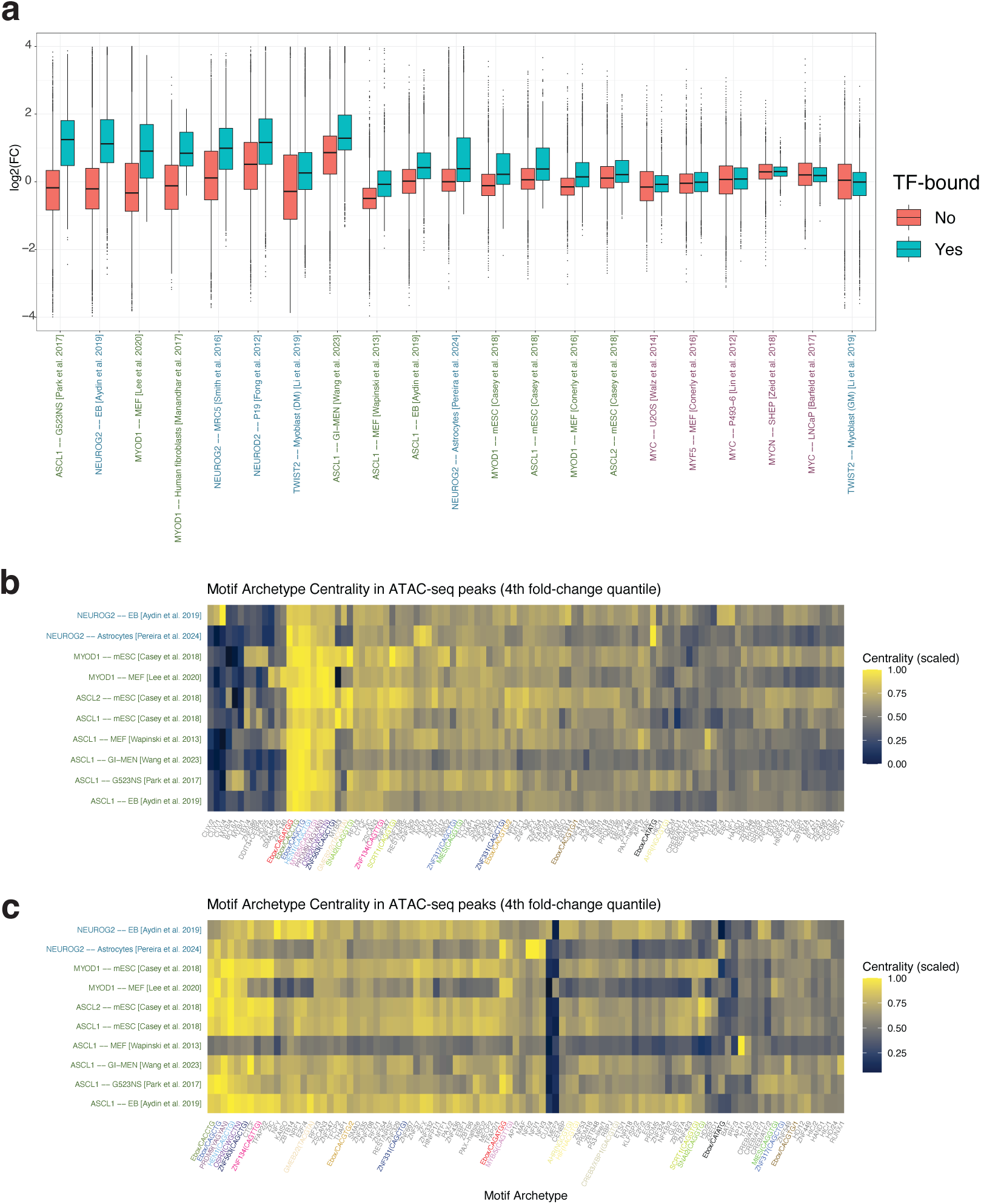
Detailed inspection of chromatin remodeling capacities of bHLH proteins and associated motif usage. **A)** Box plots displaying accessibility fold-changes of ATAC-seq and H3K27ac ChIP-seq peaks post vs pre-induction, stratified within each study based on whether they intersect with a ChIP-seq peak of the corresponding factor. Studies are ordered based on the difference between the median fold-change of the peaks that intersect vs the ones that do not intersect with the TF ChIP-seq peaks. **B)** Heatmap illustrating the centrality of Vierstra archetype motifs relative to ATAC-seq summits. Analysis is restricted to ATAC-seq peaks in the highest fold-change quartile (Q4) that specifically intersect with target transcription factor (TF) ChIP-seq peaks. The motifs shown are filtered for significance, including only archetypes ranked within the top 50 by centrality in at least one experiment. Centrality values are scaled by row (experiment). **C)** The same analysis but restricted to the ATAC-seq peaks of the 4th quartile that do not intersect with the TF ChIP-seq peaks.

**Fig. S7.**
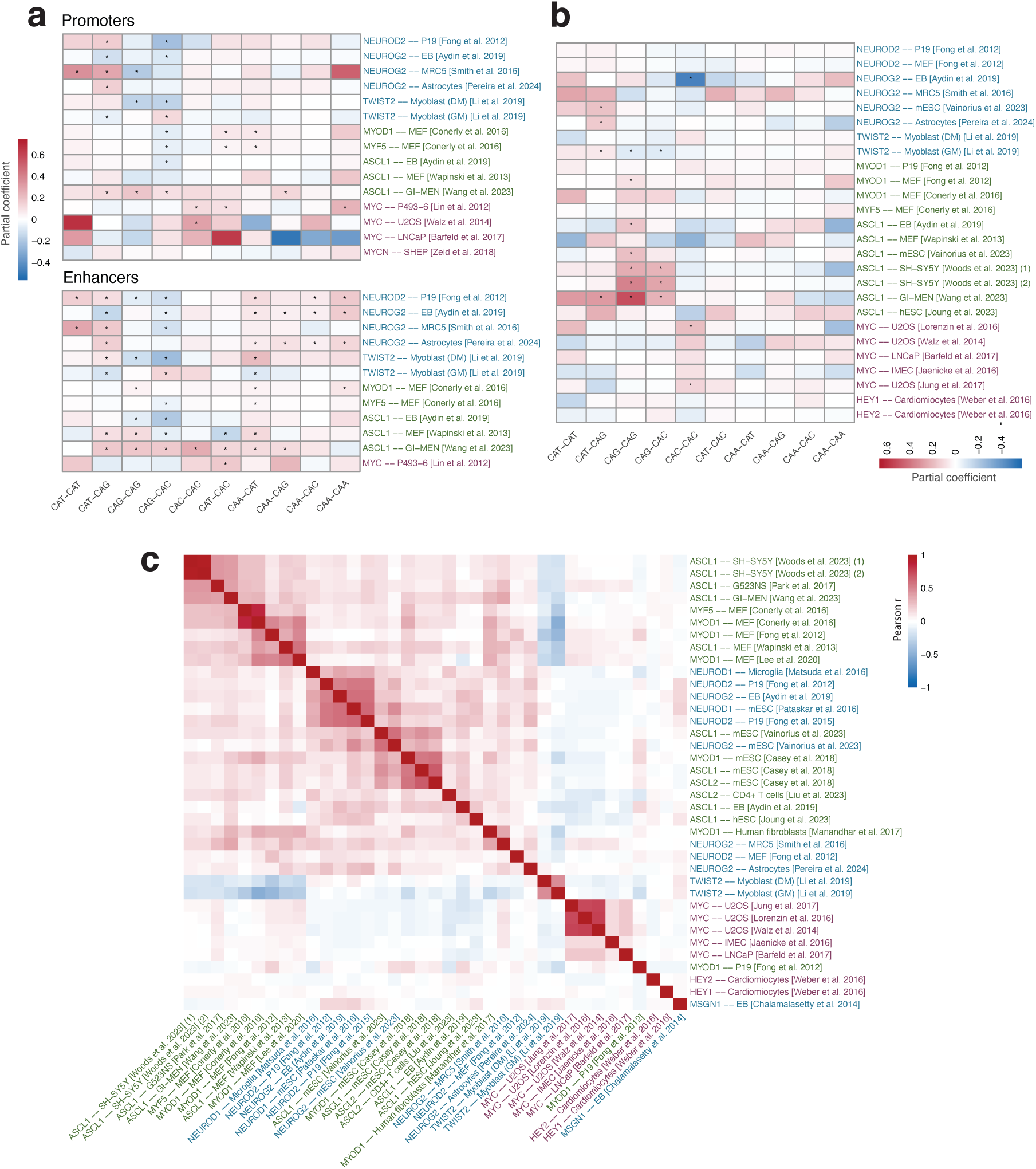
Controlled assessment of E-box-mediated transcriptional and epigenomic rewiring and cross-experiment comparison of induced transcriptomic profiles. A,B) Same as in figure 5A and B, but adding the covariate of initial chromatin accessibility. To facilitate a direct comparison of partial coefficients with and without this adjustment, the values of coefficients were mapped to the original color scales used in the previous figures. **C)** Heatmap showing Pearson correlation coefficients of gene expression fold-changes across induction experiments. Columns and rows represent individual experiments, clustered by similarity of the Pearson coefficients across studies.

**Fig. S8.**
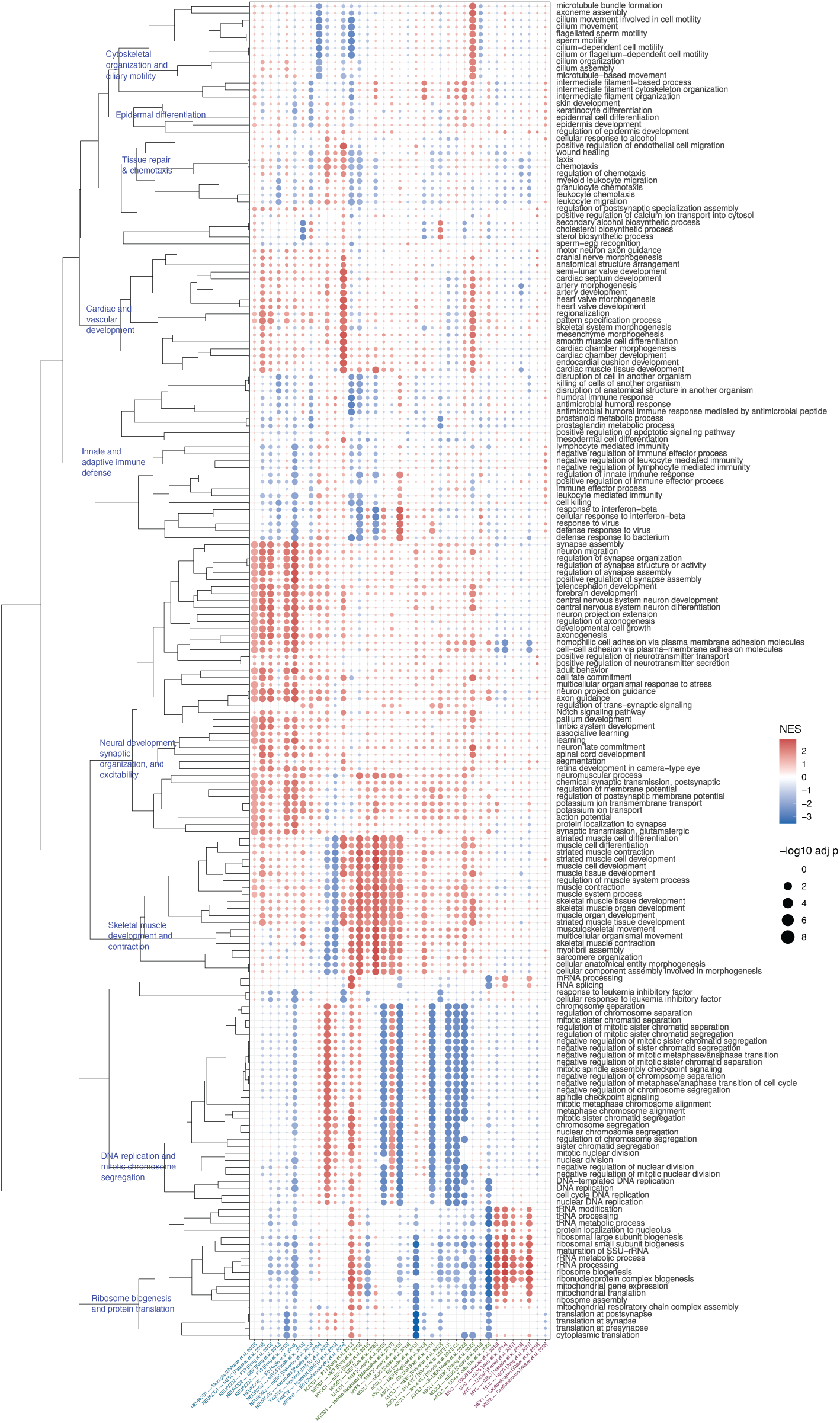
Broad description of gene ontologies. Balloon plot illustrating GSEA results for expression fold-change data. To highlight key findings, only ontologies ranking in the top 10 by significance in at least one experiment are shown. The rows are hierarchically clustered based on Normalized Enrichment Scores (NES), with the resulting dendrogram shown on the left. Major functional categories are manually annotated along the dendrogram.

